# GLP1 receptor agonism ameliorates Parkinson’s disease through modulation of neuronal insulin signalling and glial suppression

**DOI:** 10.1101/2024.02.28.582460

**Authors:** Dilan Athauda, James R Evans, Laura Mahoney-Sanchez, Gurvir S Virdi, Patricia Lopez-Garcia, Anna Wernick, Aaron Wagen, Karishma D’Sa, Joanne Lachica, Stephanie Strohbuecker, Giulia Vecchi, Craig Leighton, Rebecca S. Saleeb, Judi O’Shaughnessy, Christina E. Toomey, Nirosen Vijiaratnam, Christine Girges, Yazhou Li, Maja Mustapic, Khalida Ismail, Melanie Davies, Dimitrios Kapogiannis, Minee L Choi, Mina Ryten, Mathew H. Horrocks, Nigel Greig, Thomas Foltynie, Sonia Gandhi

## Abstract

Neuronal insulin resistance is linked to the pathogenesis of Parkinson’s disease through unclear, but potentially targetable, mechanisms. We delineated neuronal and glial mechanisms of insulin resistance and glucagon-like 1 peptide (GLP-1) receptor agonism in human iPSC models of synucleinopathy, and corroborated our findings in patient samples from a Phase 2 trial of a GLP-1R agonist in Parkinson’s (NCT01971242). Human iPSC models of synucleinopathy exhibit neuronal insulin resistance and dysfunctional insulin signalling, which is associated with inhibition of the neuroprotective Akt pathways, and increased expression of the MAPK-associated p38 and JNK stress pathways. Ultimately, this imbalance is associated with cellular stress, impaired proteostasis, accumulation of α-synuclein, and neuronal loss. The GLP-1R agonist exenatide led to restoration of insulin signalling, associated with restoration of Akt signalling and suppression of the MAPK pathways in neurons. GLP-1R agonism reverses the neuronal toxicity associated with the synucleinopathy, through reduction of oxidative stress, improved mitochondrial and lysosomal function, reduced aggregation of α-synuclein, and enhanced neuronal viability. GLP-1R agonism further suppresses synuclein induced inflammatory states in glia, leading to neuroprotection through non cell autonomous effects. In the exenatide-PD2 clinical trial, exenatide treatment was associated with clinical improvement in individuals with higher baseline MAPK expression (and thus insulin resistance). Exenatide treatment led to a reduction of α-synuclein aggregates, and a reduction in inflammatory cytokine IL-6. Taken together, our patient platform defines the mechanisms of GLP-1R action in neurons and astrocytes, identifies the population likely to benefit from GLP-1R agonism, and highlights the utility of GLP-1R agonism as a disease modifying strategy in synucleinopathies.

## Introduction

Impairment of insulin signalling in Parkinson’s disease (PD) may be related to its pathogenesis, as Type 2 diabetes (T2DM) is an established risk factor for the development of PD ^1,2^. Co-morbid T2DM in PD is associated with faster motor decline, development of cognitive impairment and depression^3–6^, which is not simply explicable by the co-occurrence of vascular pathology^4^. The hallmark of T2DM is peripheral insulin resistance (defined broadly as reduced cellular responsiveness to insulin), but the development of central, or neuronal insulin resistance (NInsR) may induce pathophysiological events in PD (Figure 1A)^7,8^. Insulin binding to its transmembrane insulin receptor (IR) triggers two major kinase cascades, the phosphoinositide 3-kinase (PI3K-Akt) and the mitogen-activated protein (MAP) kinase pathways. These modulate a variety of downstream processes including apoptosis, autophagy, inflammation, nerve cell metabolism, protein synthesis and synaptic plasticity, all of which are implicated in both PD and T2DM pathogenesis^9^ (Figure 1A).

**Figure 1.**
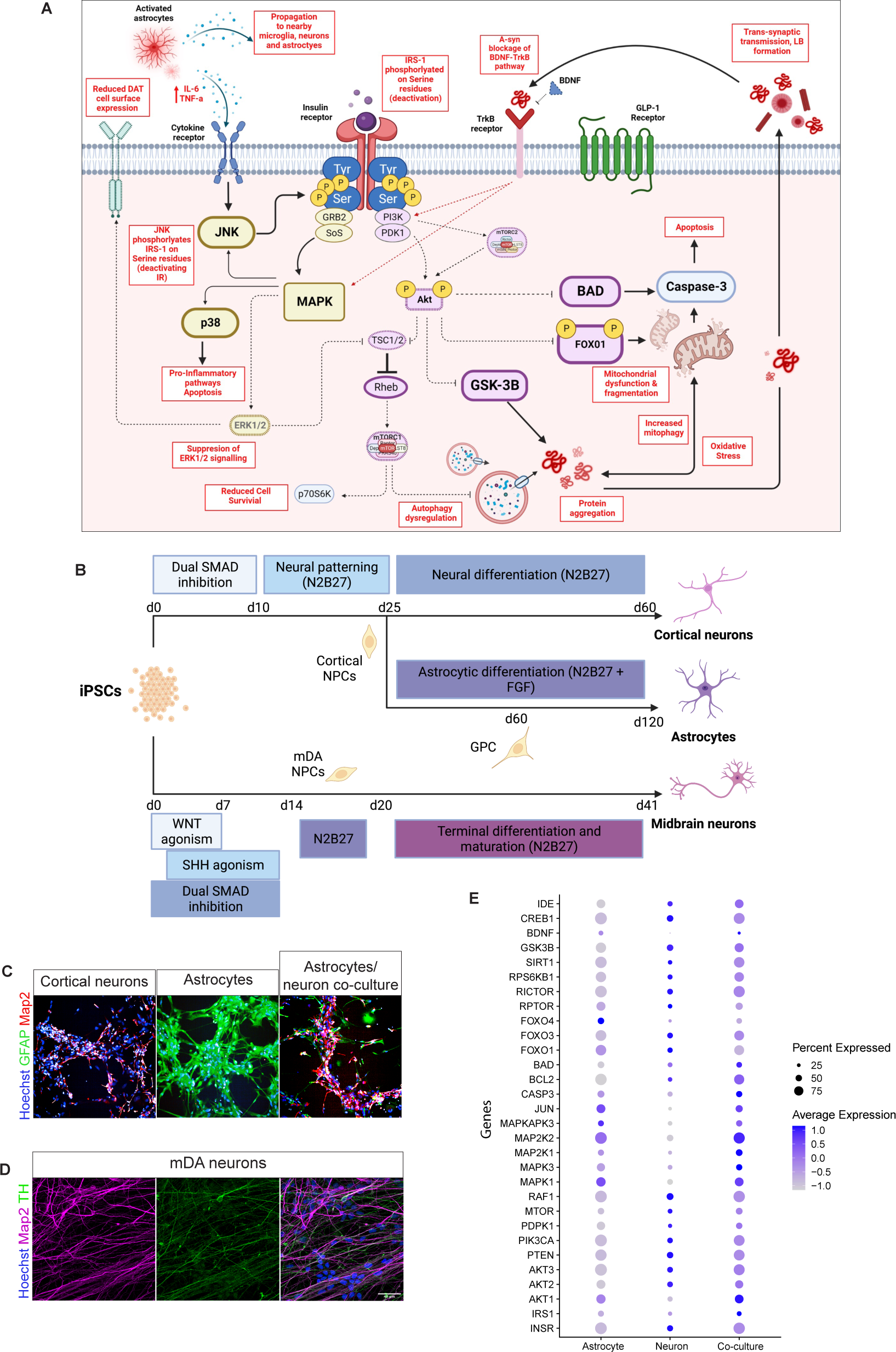
Insulin resistance signalling pathway and characterisation of cellular models used in this study. **A.** Insulin signalling pathway in neurons modulates cell survival via balanced activation of two main pathways Akt and MAPK influencing a range of cellular processes. Insulin resistance leads to serine phosphorylation of Insulin substrate-1 (IRS-1) -uncoupling of downstream insulin signalling and reduced phosphorylation of Akt. Subsequently, there is reduced inhibition of GSK-3B and TSC1/2, leading to inactivation of mTORC, which promotes protein aggregation and dysregulation of autophagy, and leads to reduced protein expression. Reduced inhibition of FOXO1 promotes oxidative stress and mitochondrial dysfunction, leading to increased oxidative stress, which further promotes a-synuclein aggregation. In parallel, less insulin stimulation leads to down regulation of ERK1/2 signalling, leading to less mTORC1 activation, and also reduced ERK-mediated DAT expression. Increased a-synuclein aggregation interferes with BDNF-TrKB signalling, further de-activating MAPK/ERK and PI3K/Akt, and also induced phosphorylation of IRS-1p on serine residues (further promoting insulin resistance). Basal SNCA astrocytes display elevated expression of IL-6 and TNF-a, further promoting stress pathways MAPK/p38 and MAPK/JNK, leading to further IRS-1p serine phosphorylation. **B.** Schematic of the differentiation protocol of human iPSCs into either cortical neurons, midbrain dopaminergic neurons (mDA) or astrocytes. **C.** Representative ICC images of cortical neurons, astrocytes and the co-culture of both astrocytes and cortical neurons. **D.** Representative ICC images of mDA neurons. **E.** Dot plot showing the expression of the markers for the insulin and GLP-1 signalling pathways observed in the single cell RNA-Seq data.

While peripheral insulin resistance may be measured by direct methods, attempts to measure NInsR “in vivo” have proven difficult^10^, though may be indirectly quantified by measuring the phosphorylation state of insulin receptor substrate-1 (IRS-1), a critical node in insulin pathway, and can enhance or diminish insulin signalling. Serine phosphorylation of IRS-1 (at residues s312/s616/s636) deactivates and decouples insulin signalling, and thus elevated expression of IRS-1 s312 is recognised as a proxy marker of NInsR^11–13^. Accordingly, elevated expression of IRS-1 s312 has been found in post-mortem studies of patients with PD^14^, Alzheimer’s Disease (AD)^10^ and Multiple System Atrophy (MSA)^15^, in association with severe dopaminergic depletion in the caudate and ventral striatum^5,16^, frontotemporal cortical atrophy^17,18^, and lower cerebral glucose hypometabolism^19^. In addition, downstream kinases of the insulin pathway are also found to be disrupted, with lower levels of phosphorylated Akt (Aktp S473) in Tyrosine hydroxylase (TH)^+^ neurons from PD patient brains compared to healthy controls^20,21^.

Glucagon-like peptide-1 receptor agonists (GLP-1RA) are “insulin-sensitizing” drugs used in the treatment of T2DM^22,23^, and activate similar downstream pathways as insulin. Interestingly, individuals with T2DM prescribed GLP-1R agonists are 36-60% less likely to develop PD compared to users of other oral antidiabetic drugs^24^, through mechanisms independent of peripheral effects^25^. Furthermore, in two clinical trials involving PD patients treated with exenatide (Ex-4) – the first-in-class GLP-1R agonist – patients in the exenatide-treated group exhibited sustained motor and non-motor benefits compared with the control/placebo groups^26,27^. Numerous clinical trials are now underway evaluating a range of GLP-1RA’s in PD to assess their effects on disease progression, but with limited insight into the underlying therapeutic mechanisms. This lack of understanding of GLP-1 RA-induced neuroprotective mechanisms in humans hinders our ability to define accurate biomarkers and optimise these promising approaches to treat PD.

We generated iPSC-derived midbrain dopaminergic and cortical neurons from healthy controls and patients with *SNCA* mutations and characterized the effect of synucleinopathy on insulin signalling in terms of insulin resistance and expression of downstream Akt and MAPK pathways. Neurons with *SNCA* mutations, and neurons exposed to α-synuclein oligomers were used to determine the effect of GLP-1R agonism on intracellular insulin signalling pathways. iPSC-derived astrocytes were used to investigate the effect of GLP-1R agonism on astrocytes. Finally, we translated these findings from *in vitro* models of synucleinopathy to biomarkers that could be measured in patient biofluids. By probing patient samples from the Exenatide-PD2 trial, we assessed the effect of exenatide on insulin signalling, inflammation, and protein aggregation states in patients.

## Results

### Generation of model

We generated cortical and midbrain dopaminergic neurons (mDA) from human iPSCs from healthy individuals and PD patients bearing *SNCA* mutations (Figure 1B). Differentiation of control iPSCs (two control independent clones), *SNCA* iPSCs (*SNCA* triplication, *A53T* x 2 lines), and two isogenic control iPSCs, into mDA neurons^24,25^, cortical neurons or cortical astrocytes^26,27^ was performed using established protocols (Figure 1B). Following terminal differentiation, the culture is highly enriched in neurons or astrocytes, displaying the cell specific markers(MAP2 for cortical neurons, TH for mDA neurons, and GFAP for astrocytes (Figure 1C & 1D)). Single-cell RNA-seq highlight the expression of the main components of the insulin and GLP-1 signalling pathway (Figure 1E) in cortical neurons and astrocytes. Data presented throughout the study represent aggregated data of the *SNCA* mutations, and the individual line by line data is presented in the Supplementary Materials(Supplementary figures 1 and 3).

### Dysregulated insulin signalling and neuronal insulin resistance occurs in *SNCA* mutant neurons

To assess the integrity of dynamic neuronal insulin signalling, day-55 post-differentiation mDA neurons were selected and cultured for a further 7 days +/-25nM exenatide, and then incubated overnight without insulin. Neurons were then treated with either 0nM, 100nM, 250nM, or 1000nM exogenous insulin for 30-minutes and cell pellets were collected for quantification of Akt phosphorylation - the activation of which reflects integrity of insulin signalling. When treated with 100nM exogenous insulin, control neurons exhibited an appropriate physiological response to insulin, producing a significant increase in pAkt of 13.8% from baseline, and with 1000nM insulin, a 35.5% increase of pAkt from baseline (Figure 2A – left panel). However, *SNCA* neurons exhibited a blunted response to increasing concentrations of insulin with no significant increases in pAkt in response to increasing exogenous insulin concentrations – akin to insulin resistance (Figure 2A – middle panel). However, *SNCA* neurons incubated with 25nM exenatide demonstrate a significant 41.4% increase in pAkt to 100nM insulin and 40.1% increase in pAkt from baseline, thus treatment with exenatide can partially restore dysfunctional insulin resistance in *SNCA* neurons (Figure 2A – right panel).

**Figure 2.**
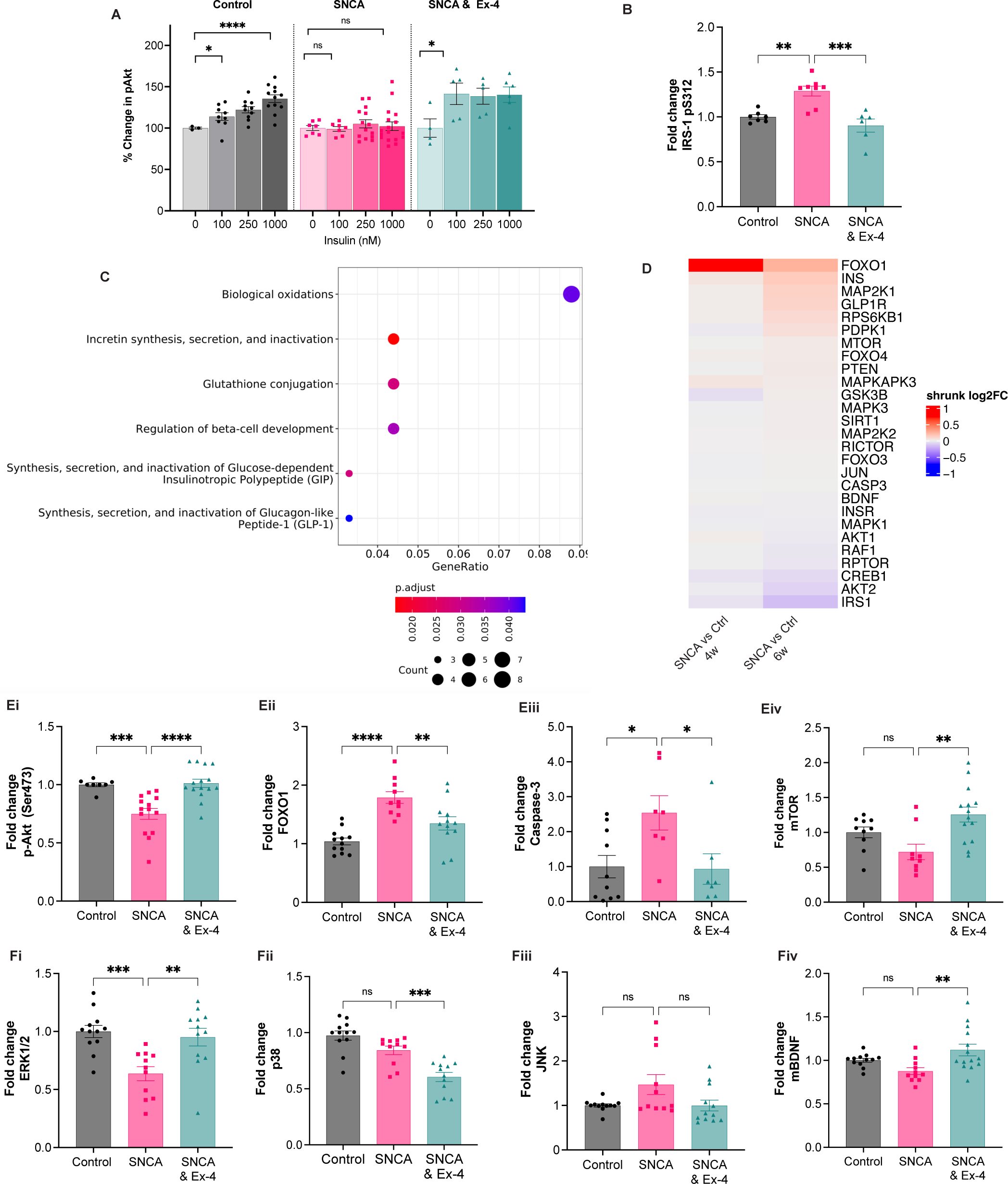
GLP-1R agonismrescues neuronal insulin resistance in human *SNCA* midbrain dopaminergic neurons. **A.** Day-55 post-differentiation mDA neurons were cultured for 7 days +/- 25nM exenatide (Ex-4), and then incubated overnight without insulin. Neurons were then treated with either 0nM, 100nM, 250nM, or 1000nM exogenous insulin for 30-minutes and cells collected for measurement of pAkt. **Left panel -** control neurons exhibit an appropriate physiological response to 100nM insulin, producing a significant increase in pAkt of 13.8% from baseline (95%CI 2.8 - 24.8, p=0.019), and with 1000nM insulin, a 35.5% increase of pAkt from baseline (95%CI 24.9 - 46.1, p=<0.0001). **Middle panel** when *SNCA* neurons were stimulated with 100nM insulin, there is no change from baseline in pAkt (-1.1%, 95%CI -10.1 – 8.0, p=0.801), and in response to 1000nM, only a 2.2% pAkt from baseline (95%CI -10.2 – 14.7, p>0.711). This is akin to insulin resistance. **Right panel** - when *SNCA* neurons were incubated with 25nM exenatide, in response to 100nM insulin, there was a significant 41.4% increase in pAkt from baseline (95%CI 0.90 – 81.9, p=0.043), while 1000nM insulin treatment caused an increase of pAkt from baseline of 40.1% (95%CI 6.3 – 74.4, p=0.025). **B.** SNCA mDA neurons demonstrate elevated basal expression of IRS-1 pS312 (a marker of insulin resistance) compared to control (CON: 1.00 ± 0.1AU: *SNCA*: 1.32 ± 0.1, p=0.0015), while SNCA neurons treated with Ex-4 had lower expression of IRS-1 pS312 (*SNCA*: 1.32 ± 0.1; *SNCA*+Ex-4: 0.90 p=0.0001) **C.** Transcriptomic data profiling showing the top 6 most differentially expressed Reactome pathways between *SNCA* and control mDA neurons at 4 weeks, of which 3 pathways involve downregulation of GLP-1 / incretin synthesis **D.** Based on the differential gene expression analysis between *SNCA* and control samples at each of the investigated time points during the differentiation, regularised log2 fold changes are visualised for a subset of genes selected based on the GLP-1 and insulin signalling pathways. **E.** Compared to control neurons, SNCA neurons dsisplayed dysfunctional downstream insulin signalling with (**Ei**), reduced expression of pAkt S473 (CON: 1.0 ± 0.1AU: *SNCA*: 0.75 ± 0.1AU, *SNCA*+Ex-4: 1.01 ± 0.1AU, p=0.0007), (**Eii**) increased FOXO1 (CON: 1.0 ± 0.1AU: *SNCA*: 1.79 ± 0.1AU, *SNCA*+Ex-4: 1.35 ± 0.1AU, p<0.0001) and (**Eiii**) increased caspase-3 expression (CON: 1.0 ± 0.3AU: *SNCA*: 2.53 ± 0.3AU, *SNCA*+Ex-4: 0.93 ± 0.1AU p=0.0212), and (**Eiv**) reduced mTOR levels (CON: 1.0 ± 0.2AU, *SNCA*: 0.72 ± 0.3AU, *SNCA*+Ex-4: 1.26 ± 0.4AU p=0.0041), all pf which were reversed with Exenatide treatment **F.** Compared to control neurons, SNCA neurons also have dysfunctional MAPK signalling and showed (**Fi**) reduced ERK1/2 (CON: 1.0 ± 0.1AU: *SNCA*: 0.63 ± 0.1AU, *SNCA*+Ex-4: 0.95 ± 0.1AU p=0.0007), however there was no difference (**Fii**) in p38 (CON: 1.0 ± 0.1AU: *SNCA*: 0.84 ± 0.1AU, *SNCA*+Ex-4: 0.60 ± 0.1AU p<0.0001), (**Fiii**) p-JNK (CON: 1.0 ± 0.1AU: *SNCA*: 1.46 ± 0.2AU, *SNCA*+Ex-4: 0.99 ± 0.1AU p=0.0435) but (**Fiv**) exenatide did significantly increase mBDNF signal in SNCA neurons (CON: 1.0 ± 0.1AU*, SNCA*: 0.87 ± 0.1AU, *SNCA*+Ex-4: 1.11 ± 0.2AU p=0.0031) Note: Statistical significance measured by one-way ANOVA with Bonferroni multiple comparisons test. Fold change in mean absorbance signal measured relative to control. Each data point represents analyte quantification from an individual well of cultured neurons. *SNCA* group comprise data pooled from A53T & SNCAx3 mutations (see Supplementary data for data split by *SNCA* mutation). Data represented as mean (normalised to control) ± SEM of at least three independent experiments, *p < 0.05, **p < 0.005, ***p < 0.0005, ****p < 0.0001.

In a separate experiment, neurons were cultured until day-55 post-differentiation, then cultured for a further 7 days +/- 25nM exenatide treatment to assess basal markers of insulin resistance by measuring the basal expression of IRS-1 pS312. Expression of IRS-1p S312 was significantly higher in *SNCA* mDA neurons compared to controls at basal conditions. However, exenatide-treated neurons had significantly reduced IRS-1p S312 expression compared to untreated *SNCA* (Figure 2B). Thus, both dynamic and basal characterisation demonstrates insulin resistance in *SNCA* neurons that is reversed by treatment with exenatide.

### *SNCA* mutant neurons display altered incretin pathways, Akt and ERK 1/2 signalling pathways

We performed bulk RNA-sequencing comparing the transcriptomes of *SNCA* (2 x A53T lines, 1 x *SNCAx3*) versus healthy control mDA neuronal cultures at 4 and 6 weeks of differentiation.

At four weeks of differentiation, this yielded 316 significantly (FDR<0.05) differentially expressed genes (DEGs). Subsequently, we identified six over-represented Reactome pathways in the set of 316 DEGs(Figure 2C), which included pathways involved in oxidative stress (commonly known to be altered in PD pathogenesis): dysregulation of biological oxidation; glutathione conjugation, and regulation of beta cell development. However, 3 pathways were involved in incretin (GLP-1 or glucose-dependent insulinotropic polypeptide (GIP)) synthesis, which act to stimulate insulin secretion – thus identifying the incretin pathway and insulin secretion as a potential target in altering disease pathogenesis. While incretin downregulation has been shown to occur peripherally in patients with Type 2 DM^32^, this downregulation of the incretin pathway has not previously been observed in neurons. The underlying individual genes comprising these dysregulated Reactome pathways are shown in Supplementary Figure 2.

We next identified gene marker sets based on the GLP-1 and insulin signalling pathway and observed a subset of the members of the GLP-1 and insulin signalling pathway are differentially expressed between *SNCA* ( 2 x A53T lines, 1 x SNCA) and control (Ctrl1, Ctrl 3, Ctrl4) neurons: IRS-1, Akt, CREB, GSK3B, RPTOR are downregulated, while FOXO1, INS, MAPK, GLP-1R, RPS6KB1 are upregulated in comparison to control neurons (Figure 2D). Given that MAPK and RPS6KB1 are known to be increased in insulin resistance, this supports our previous data implying the presence of dysfunctional insulin signalling in *SNCA* neurons.

We quantified protein expression of key kinases in the Akt and the MAPK signalling pathways (the primary effectors that reflect intact insulin signalling) in *SNCA* and control neurons using ELISA based assays. Under basal conditions, *SNCA* mDA neurons exhibited significantly altered signalling of both Akt and MAPK pathways (Figure 2 E&F) compared to control neurons. *SNCA* neurons displayed a significant reduction in expression of p-Akt S473 (Figure 2Ei). In healthy neurons, Akt activation typically inhibits forkhead transcriptional factor O1 (FOXO1) and Caspase-3, two kinases involved in promoting cell-death mechanisms. FOXO1 is a critical regulator of metabolic homoeostasis and acts as a pro-apoptotic regulator ^33^ and inhibitor of TH gene transcription, resulting in decreased TH expression^34^; whereas Capase-3 is a primary mediator of cell death. Correspondingly, we observed *SNCA* neurons exhibit significantly elevated expression of phosphorylated FOXO1 (Figure 2Eii), and cleaved Caspase-3 (Figure 2Eiii).

The MAPK pathway is the other pathway modulated by insulin signalling, and can be simplified into three main branches - JNKs, the p38 kinases and the ERKs. JNK and p38 activation are typically thought to be involved in responding to and promoting oxidative stress and inflammation, whereas ERKs respond to growth factors to promote cell growth and differentiation^35^. At baseline, *SNCA* neurons had significantly reduced expression of ERK1/2 compared to controls (Figure 2Fi); but there was no significant differences in the basal levels of the other MAPK kinases p38 (Figure 2Fii) and p-JNK (Figure 2Fiii).

Taken together, these results indicate dysfunctional / disrupted insulin signalling in *SNCA* mutant neurons compared to healthy controls, which is associated with promotion of cell-death pathways.

### GLP-1R agonism restores Akt and downstream MAPK signalling pathways in *SNCA* neurons

We utilized the GLP-1R agonist, exenatide, to test the modulation of neuronal insulin resistance, and its downstream consequences on insulin signalling. Neurons were cultured until day-55 post-differentiation, then cultured for a further 7 days +/-25nM exenatide treatment and cells were collected for ELISA assays. *SNCA* neurons treated with exenatide demonstrated significant increases in p-Akt S473 (Figure 2 Ei); with concomitant significant reductions in FOXO1 (Figure 2Eii), and cleaved Caspase-3 (Figure 2Eiii), and significantly increased mTOR expression (Figure 2Eiv) compared to untreated *SNCA* neurons. Furthermore, exenatide-treated *SNCA* neurons had significantly increased ERK1/2 signalling (Figure 2Fi), and suppression of p38 (Figure 2Fii), and p-JNK expression (Figure 2Fiii).

BDNF/TrkB is highly expressed and activated in the dopaminergic neurons of the substantia nigra and plays a critical role in the maintenance of neurons^36^; and expression of BDNF/TrkB in the substantia nigra is significantly reduced in studies of PD patients compared to controls^37^. Correspondingly, exenatide significantly elevated mBDNF expression compared to untreated *SNCA* neurons (Figure 2Fiv).

In summary, under basal conditions, mDA *SNCA* neurons display dysregulated intracellular insulin signalling, with reduced Akt phosphorylation and ERK1/2 signalling, leading to increased activation of FOXO1 and promotion of cell death pathways. Treatment with exenatide preserves Akt and ERK1/2 expression, suppresses stress pathways JNK and p38, and may restore mTORC1 function - enhancing protective /cell survival pathways.

### GLP-1 R agonism improves cellular stress, and prevents neurotoxicity in *SNCA* neurons

We previously reported cellular pathology and toxicity associated with the *SNCA* triplication (*SNCA*x3) and A53T mutation in mDA neurons^28,29,38–40^. We quantified cell viability using the dye sytox green, which is excluded from healthy cells (Figure 3A). There was a significant increase in cell death in *SNCA* neurons compared to control neurons, however, exenatide treatment effectively preserved cellular viability in *SNCA* neurons similar to controls (Figure 3B). α-synuclein aggregates were labelled with a p129 α-synuclein antibody in day 65 *SNCA* mDA neurons, which demonstrated *SNCA* neurons exhibited a significant increase in phosphorylated aggregates of α-synuclein compared to controls, significantly reduced by exenatide treatment (Figure 3C).

**Figure 3.**
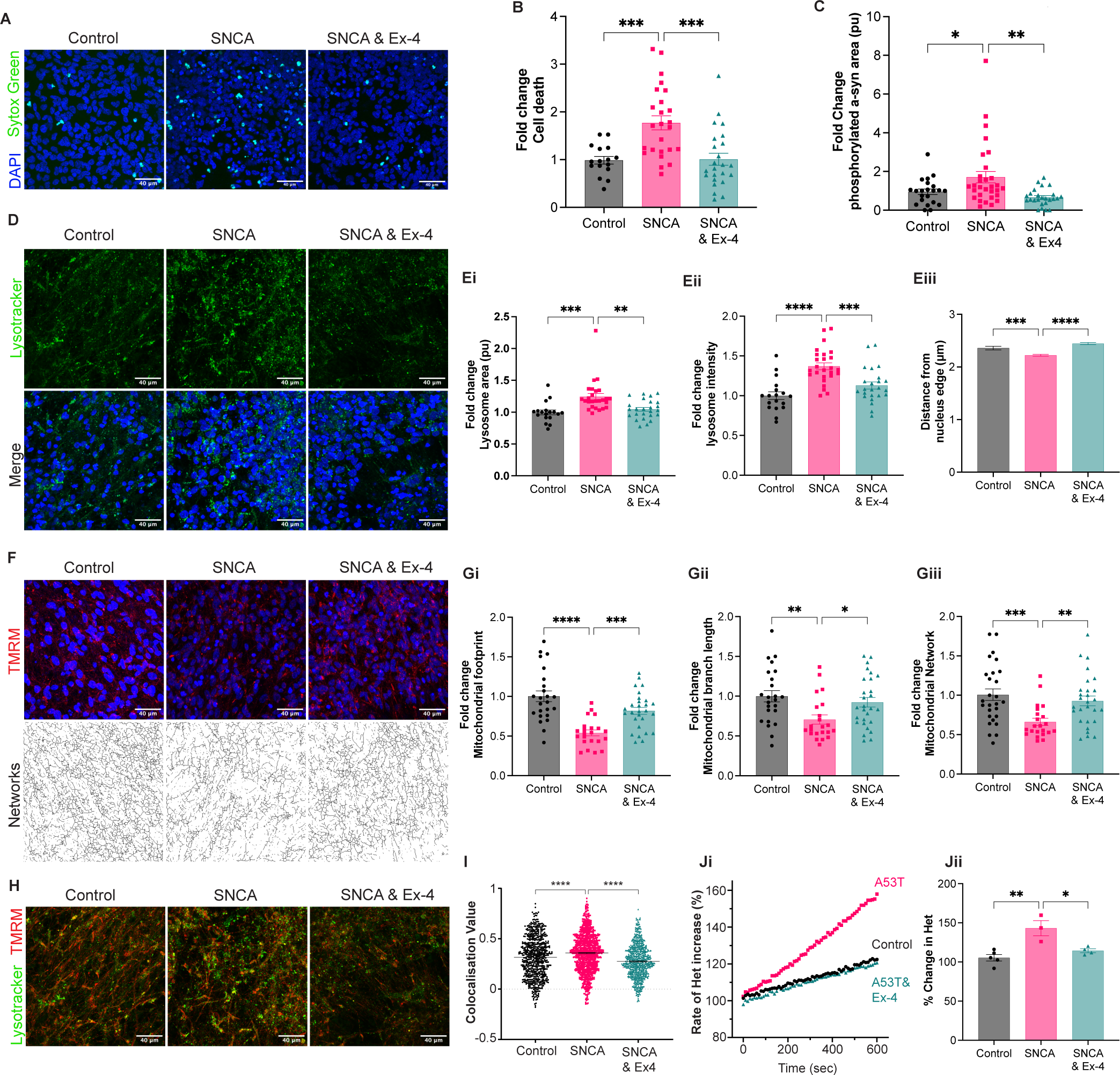
GLP-1 R agonism improves cellular stress, and prevents neurotoxicity in *SNCA* neurons. **A.** Representative live-cell images of day 65 mDA neurons (Control, *SNCA* and *SNCA* neurons treated with Exenatide (SNCA & Ex-4) depicting dead cells labelled with Sytox green. **B**. There was a significant increase in cell death in *SNCA* neurons compared to control neurons, however, exenatide treatment effectively preserved cellular viability in *SNCA* neurons similar to controls (CON:1.0 ± 0.1%; *SNCA* 1.77 ± 0.1, *SNCA*+Ex-4: 1.03, p=0.0001 **C.** Quantification of the fold change of average puncta size of α-synuclein aggregates demonstrated *SNCA* neurons exhibited a significant increase in phosphorylated aggregates of α-synuclein compared to controls, but exenatide treatment significantly reduced phosphorylated aggregates of α-synuclein (CON: 1.0 ± 0.1; *SNCA*: 1.71 ± 0.2, *SNCA* +Ex-4: 0.66 ± 0.1, p=0.0023). **D.** Representative live-cell images of lysosome using Lysotracker Deep Red (green dye) and merged images with Hoechst nuclear stain (blue dye) in day 65 of differentiation mDA neurons. Quantification of the fold change in lysosomal area **(Ei)** SNCA neurons display increased lysosomal area (CON: 1.0 ± 0.1: *SNCA*: 1.25 ± 0.1, *SNCA*+Ex-4 1.05 ± 0.1p=0.0001), **(Eii)** increased integrated intensity (CON: 1.0 ± 0.1; *SNCA* 1.37 ± 0.2, *SNCA* +Ex-4: 1.12 ± 0.1, p<0.0001) and **(Eiii)** increased distance from the nuclear edge (CON: 2.3 ± 0.03nm; *SNCA*: 2.22 ± 0.01px; *SNCA*+Ex-4: 2.45 ± 0.01px, p<0.0001) – suggesting overactivation / stress of this pathway, while exenatide treatment significantly attenuates these effects **F.** Representative live-cell images showing reduced mitochondrial membrane potential in *SNCA* mDA as measured by the lipophilic cationic dye TMRM (red dye), and skeletonized images of mitochondrial networks used to analyse integrity/fragmentation of mitochondrial networks. Foldchange of mitochondrial footprint **(Gi)** We utilised the MiNA toolset to analyse mitochondrial integrity, and demonstrated *SNCA* neurons display reduced mitochondria footprint (area occupied by mitochondria normalised to cell number) (CON:1.0 ± 0.1; *SNCA*: 0.53 ± 0.1, *SNCA*+Ex-4: 0.82 ± 0.1, p=0.0005)**, (Gii)** mean branch length “length of all rods/branch mitochondrial structures” (CON: 1.0 ± 0.1; *SNCA*: 0.70 ± 0.1, *SNCA*+Ex-4: 0.92 ± 0.1, p=0.0359) and **(Giii)** network size (CON: 0.1 ± 0.1; *SNCA*: 0.66 ± 0.1, *SNCA* +Ex-4: 0.93 ±0.1 , p=0.0004) – indicating mitochondrial dysfunction – all attenuated by Exenatide treatment. **H.** Representative live-cell images of day 65 mDA neurons stained with TMRM (red - mitochondria) and Lysotracker DeepRed (green - lysosomes) to act as a proxy marker for mitophagy. I. Quantification of the co-localization of mitochondria and lysosomes was significantly higher in *SNCA* neurons compared to controls, whereas treatment with exenatide prevented the increase in co-localization (CON: 31.5 ± 0.8%; *SNCA*: 35.8 ± 0.6%, *SNCA*: 35.8 ± 0.6%; *SNCA* +Ex-4: 27.5 ± 0.5%, p<0.0001). **Ji.** Trace showing the ratiometric measurement of superoxide generation over time measured with dihydroethidium (Het) indicating SNCA neurons have a higher basal rate of production of superoxide generation (normalised by exenatide)**. Jii.** Percentage change in the rate of superoxide production in *SNCA* neurons was significantly higher compared to controls *attenuated by exenatide) CON:106.5 ± 3.8%; *SNCA*:143.2 ± 9.7%; *SNCA* +EX4: 114.3 ± 2.4%; p=0.0111). Note: Images analysed using CellProfiler software. Statistical significance measured by One-way ANOVA Bonferroni multiple comparisons test. *SNCA* group comprise data pooled from A53T & SNCAx3 mutations (see Supplementary data for data split by *SNCA* mutation. Data represented as mean ± SEM of at least three independent experiments, *p < 0.05, **p < 0.005, ***p < 0.0005, ****p < 0.0001, Scale bar = 40um

We measured the lysosomal compartment in neurons using LysoTracker Deep red; a cationic fluorescent dye that accumulates in acidic cellular compartments (Figure 3D). Compared to healthy control neurons *SNCA* neurons demonstrated a significant increase in both the size of lysosomes (Figure 3Ei) and increase in fluorescent intensity (Figure 3Eii). Autophagy is associated with peri-nuclear clustering of lysosomes, and we observed lysosomes in the *SNCA* neurons localized closer to the perinuclear area (Figure 3Eiii). Treatment of *SNCA* neurons with exenatide resulted in restoration in both lysosome size, and intensity, and lysosomes located further away from the peri-nuclear zone (Figure 3Ei-iii).

We assessed mitochondrial function of neurons using the lipophilic cationic dye tetramethylrhodamine methyl ester (TMRM), to measure Δψ_m_, and utilized a mitochondrial network morphology analysis (MiNA) tool^41,42^ to analyze integrity of mitochondrial networks (Figure 3F). *SNCA* neurons demonstrated significant reductions in “mitochondrial footprint” (the area occupied by mitochondrial structures) (Figure 3Gi), reduction in mean “branch length” (the mean length of all rods/branch mitochondrial structures” (Figure Gii), and significant reduction in “network size” (mitochondrial structures with >1 node and three branches) (Figure 3Giii) – suggesting greater fragmentation of mitochondria in *SNCA* neurons. However, exenatide treated *SNCA* neurons exhibited preserved mitochondrial footprint, mean branch length, and mitochondrial network size compared to untreated *SNCA* neurons.

Next, neurons were co-loaded with LysoTracker Deep Red and TMRM fluorescent dyes to concurrently label lysosome and mitochondria, respectively, to allow identification of polarised mitochondria colocalized with lysosomes as an indicator of mitochondrial clearance (Figure 3H). The co-localization value (measured by colocalization and correlation between intensities in different colour channels on a pixel-by-pixel basis, within identified objects of mitochondria and lysosomes) was significantly higher in *SNCA* neurons compared to controls, whereas treatment with exenatide prevented the increase in co-localization (Figure 3I).

We measured the generation of cytosolic reactive oxygen species (ROS) using the fluorescent reporter dihydroethidium (HEt). The rate of superoxide production in *SNCA* neurons was significantly higher compared to controls but exenatide treatment significantly reduced *SNCA*-induced ROS generation (Figure 3Ji & Jii).

Exenatide protects neurons from proteotoxic stress via its action on the GLP-1R.

Oligomeric α-synuclein induces mitochondrial dysfunction, oxidative stress, ferroptosis and cell toxicity^29,38–40^ and we tested whether exenatide could prevent oligomer-induced toxicity via the GLP-1R. Control cortical neurons were pre-treated with either exenatide (100nM) alone, or exenatide + the GLP-1R Antagonist Ex9-39 (1µM), followed by 1µM oligomeric (consisting of 1µM monomer, and 10nM oligomer) α-synuclein treatment for 24 hours.

Application of α-synuclein oligomers led to an increase in cell death, which was prevented in neurons pre-treated with exenatide. These protective effects were lost in the presence of GLP-1R blockade by the GLP-1R antagonist Ex9-39 (Figure 4A & B).

**Figure 4.**
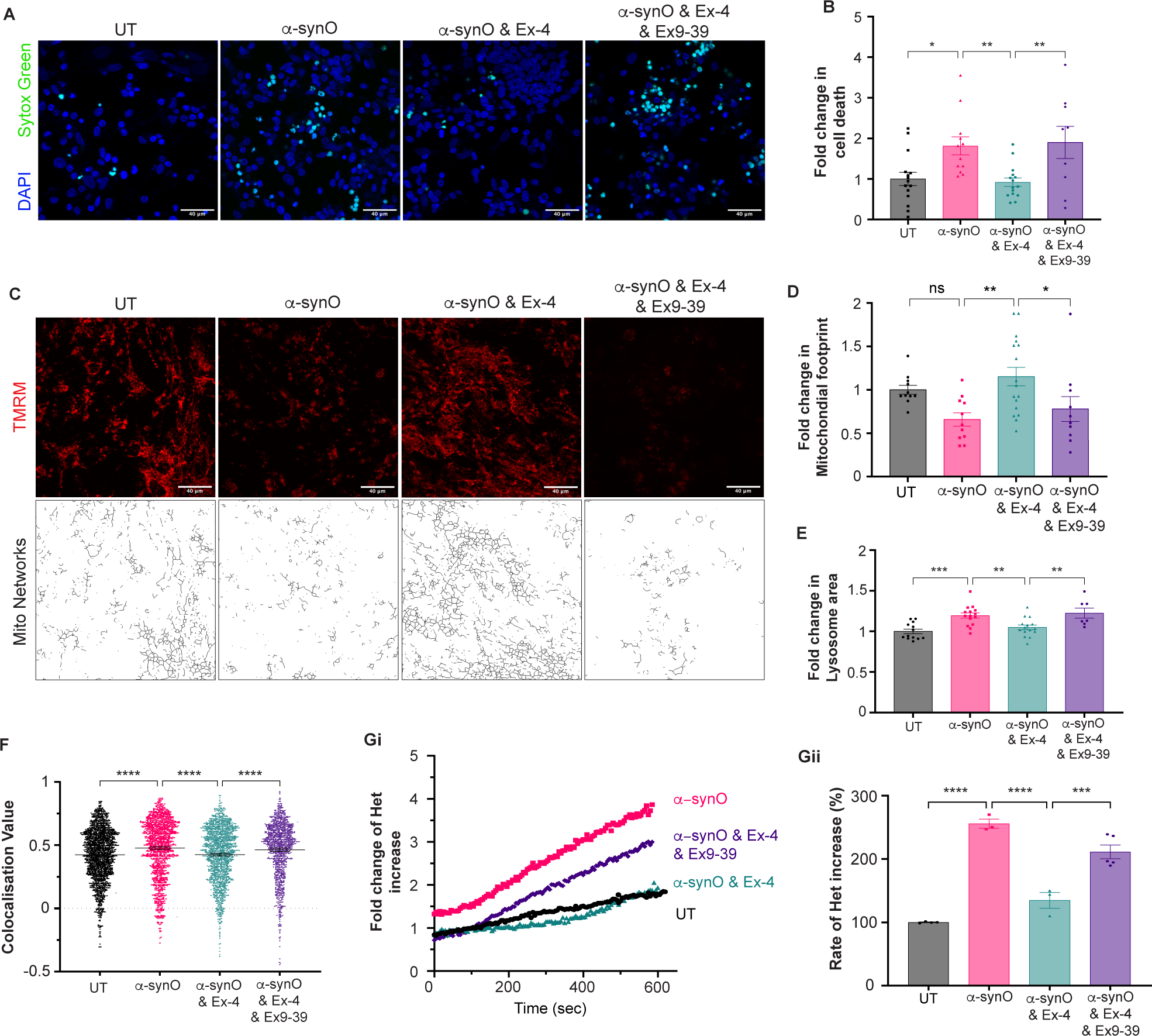
GLP-1R agonist reverses cellular pathology and neurotoxicity induced by α-synuclein oligomer treatment. **A.** Representative live-cell images of cortical neurons depicting dead cells labelled with SYTOX green in untreated neurons (UT), neurons treated overnight with α-synuclein oligomers (α-synO), neurons pre-treated with Exenatide and overnight with α-synuclein oligomers (α-synO & Ex-4), and neurons pre-treated with Exenatide, a GLP-1 antagonist (Ex9-39) and overnight with α-synuclein oligomers (α-synO & Ex-4 & Ex-9-39). **B.** Application of α-synO induced an increase in cell death, which was prevented in neurons pre-treated with exenatide, but not in the presence of the GLP-1R antagonist (CON:1.0 ± 0.1; CON+ α-synO: 1.81 ± 0.1, CON+ α-synO +EX4: 0.92 ± 0.1, CON+ α-synO +EX4+EX9-39: 1.90 ± 0.1 p=0.0015). **C.** Representative live-cell images of control cortical neurons ± α-synO ± Ex-4 ± Ex9-39 stained with TMRM and skeletonized images of mitochondrial networks and Lysotracker DeepRed. **D.** α-synO caused a significant reduction in mitochondrial footprint. However, when pre-treated with exenatide, there was a significant increase in the mitochondrial footprint compared to α-synO treated neurons, which was lost in the presence of the GLP1-R Antagonist Ex9-39 - CON: 1.0 ± 0.1, CON+ α-synO: 0.68 ± 0.1, CON+ α-synO +EX14: 1.25 ± 0.1, CON+ α-synO +EX4+GLP-1RA: 0.72 ± 0.1, p=0.0049) **E.** Application of α-synO was associated with a significant increase in lysosomal area, prevented by pre-treatment with exenatide – but not by treatment with GLP1-R Antagonist Ex9-39, (CON:1.0 ± 0.1, CON+ α-synO:1.19 ± 0.1; CON+ α-synO +EX4: 1.04 ± 0.1; CON+ α-synO +EX4+EX9-39: 1.22 ± 0.1, p<0.0001). **F**. Quantification of the co-localization of TMRM and lysotracker was significantly higher in neurons treated with oligomers compared to controls, and pre-treatment with exenatide prevented an increase in co-localization whilst pre-treatment along with the GLP1-R Antagonist Ex9-39 had no effect (CON: 42.3 ± 0.01%; CON+ α-synO: 47.6 ± 0.01%; CON+ α-synO +Ex4: 42.5 ± 0.01%; CON+ α-synO +Ex4+ EX9-39: 46.3 ± 0.01%, p<0.0001). **Gi.** Trace showing the ratiometric measurement of superoxide generation over time measured with dihydroethidium (Het). **Gii.** Application of α-synuclein oligomers to cortical neurons significantly increased ROS by 155%, and exenatide pre-treatment prevented oligomer-induced ROS generation, but pre-treatment with GLP1-R Antagonist Ex9-39 did not (CON:100 ± 0.6%; CON+ α-synO 255.7 ± 7.2%; CON+ α-synO +Ex4: 134.6 ± 12.5%; CON+ α-synO +Ex4+ EX9-39: 211.2 ± 10.8%, p<0.0001). Note: Images analysed using CellProfiler software. Statistical significance measured by One-way ANOVA Bonferroni multiple comparisons test. *SNCA* group comprise data pooled from A53T & SNCAx3 mutations (see Supplementary data for data split by *SNCA* mutation. Data represented as mean ± SEM of at least three independent experiments, *p < 0.05, **p < 0.005, ***p < 0.0005, ****p < 0.0001, Scale bar = 40um

Oligomeric α-synuclein has been shown to impair mitochondrial dysfunction, leading to fragmentation of mitochondrial networks. We utilized the MiNA tool to quantify mitochondrial networks which demonstrated oligomers caused a significant reduction in mitochondrial footprint. However, when pre-treated with exenatide, there was a significant increase in the mitochondrial footprint compared to oligomer treated neurons, which was lost in the presence of the GLP-1R Antagonist Ex9-39 (Figure 4C&D). Application of α-synuclein oligomers was also associated with a significant increase in lysosomal area, prevented by pre-treatment with exenatide – but not by treatment with GLP-1R Antagonist Ex9-39 (Figure 4E). Co-localization of TMRM and lysotracker was significantly higher in neurons treated with oligomers compared to controls and pre-treatment with exenatide prevented an increase in co-localization, whilst pre-treatment along with the GLP-1R Antagonist Ex9-39 had no effect (Figure 4F).

Application of α-synuclein oligomers to cortical neurons significantly increased ROS by 155% and exenatide pre-treatment prevented oligomer-induced ROS generation, but pre-treatment with GLP 1R Antagonist Ex9-39 blocked the restorative effect of exenatide (Figure 4Gi &Gii).

### Exenatide dampens inflammatory activation of astrocytes via the GLP-1R receptor

We evaluated the effect of GLP-1R agonism on astrocytic activation, and subsequently, neuronal health. Control astrocytes were incubated for 24hrs with either untreated astrocyte-media +/- α- synuclein oligomers (1µM) (α-syn O) +/- exenatide 25nM +/- GLP-1R Antagonist Ex9-39. Subsequently the astrocyte-conditioned media (ACM) was then incubated with neuronal cultures for 24 hours and neuronal cell viability was quantified using the dye sytox green (Figure 5A&B). Neurons incubated with α-synO-ACM had significantly increased neuronal cell death compared to neurons incubated with media from untreated astrocytes; however, neurons incubated with media from astrocytes pre-treated with exenatide were protected from the α-synO-ACM-mediated induced neuronal death, such that the neuronal cell viability was indistinguishable from untreated. The protective effects of exenatide were lost by co-treatment with the GLP-1R antagonist Ex9-39 (Figure 5B).

**Figure 5.**
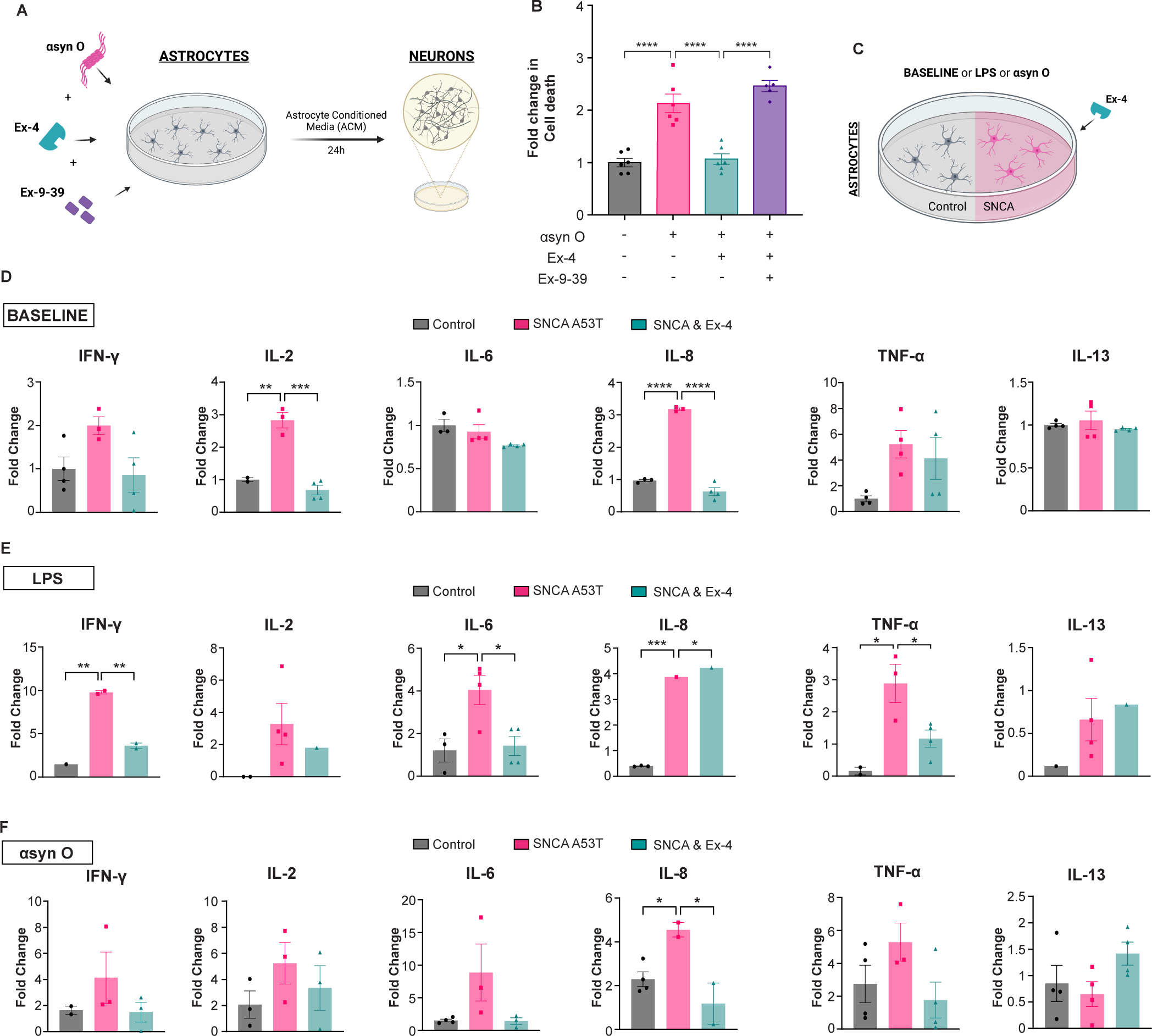
Exenatide dampens inflammatory activation of astrocytes via the GLP-1R receptor. **A.** Schematic representation of the experimental paradigm where a multiplex cytokine assay was used to measure cytokine expression from the conditioned media of control or *SNCA* astrocytes under either basal conditions (BASELINE), and then overnight treatment with Lipopolysaccharides (LPS), or treatment with α-synuclein oligomer (α-synO) +/- Exenatide (Ex-4) 25nM +/- GLP1-R Antagonist Ex9-39. **B.** Astrocyte-conditioned media (ACM) was then incubated with neuronal cultures for 24 hours and neuronal cell viability was quantified using the dye sytox green **C.** Neurons incubated with α-synO-ACM had increased neuronal cell death compared to neurons incubated with media from untreated astrocytes, however neurons incubated with media from astrocytes pre-treated with exenatide were protected from this astrocyte-mediated induced neuronal death, though the protective effects of exenatide were lost by co-treatment with the GLP-1R antagonist Ex9-39 (CON:1.0 ± 0.01; α-synO-ACM: 2.13 ± 0.1; α-synO-ACM + Ex-4: 1.06 ± 0.1; α-synO-ACM+Ex-4+Ex9-39: 2.46 ± 0.1, p<0.0001). **D.** Under basal conditions, *SNCA* astrocytes express higher levels of inflammatory cytokines IL-2 (α-synO CON:1.0 ± 0.2; *SNCA*: 2.83 ± 0.2, *SNCA*+Ex-4: 0.68 ± 0.2, p<0.0001) and IL-8 (CON:1.0 ± 0.2; *SNCA*: 3.17 ± 0.2, *SNCA*+Ex-4: 0.63 ± 0.2, p<0.0001) compared to control astrocytes, which was attenuated by Ex-4 treatment. **E.** In response to LPS exposure, SNCA astrocytes demonstrate significantly greater activation of pro-inflammatory cytokines **Ei** IFN-y, (CON:1.47; *SNCA*: 9.77 ± 0.2, *SNCA*+Ex-4: 3.61 ± 0.2, p=0.0043), **Eiii** IL-6 (CON:1.2 ± 0.2; *SNCA*: 4.05 ± 0.2, *SNCA*+Ex-4: 1.42 ± 0.2, p<0.0137), **Eiv** IL-8 (CON:0.3 ± 0.01; *SNCA*: 3.83 ± 0.2, *SNCA*+Ex-4: 4.3 ± 0.2, p<0.0001), **Ev** TNF- α (CON:0.15 ± 0.2; *SNCA*: 2.83 ± 0.2, *SNCA*+Ex-4: 1.16 ± 0.2, p<0.0132); and **Fiv** after overnight exposure to α-synO, had elevated IL-8 (CON:2.28 ± 0.2; *SNCA*: 4.45 ± 0.2, *SNCA*+Ex-4: 1.17 ± 0.2, p<0.0132) – but all attenuated with exenatide demonstrating a consistent anti-inflammatory effect in all conditions. Note: Images analysed using CellProfiler software. Statistical significance measured by One-way ANOVA with Bonferroni multiple comparisons test, *p < 0.05, **p < 0.005, ***p < 0.0005, ****p < 0.0001.

Media was collected from control and *SNCA* A53T astrocytes pre-treatment and post-treatment (baseline, α-syn O (1µM) or lipopolysaccharide (LPS) +/- exenatide 25nM) and used to quantify cytokine expression by a multiplex MSD electrochemiluminescence assay (V-PLEX Proinflammatory Panel 1) (Figure 5C).

Under basal conditions, astrocytes derived from patients with *SNCA* A53T mutation secrete significantly higher levels of pro-inflammatory cytokines IL-2 and IL-8 (Figure 5D). Following incubation with LPS, *SNCA* astrocytes demonstrated a significantly greater pro-inflammatory response compared to control astrocytes secreting greater IFN-y, IL-2, IL-6, IL-8, TNF-a in comparison to control astrocytes (Figure 5E). In addition, *SNCA* astrocytes displayed a significantly exaggerated response to α-synuclein oligomers compared to control astrocytes, secreting significantly higher levels of IL-8 compared to control neurons (Figure 5F). However, exenatide pre-treatment effectively attenuated the inflammatory response induced by α-synuclein, reducing secretion of IL-8 – with similar trends in reducing IFN-y, IL-2, IL-6, TNF-a.

### Insulin receptor signalling is altered in sporadic Parkinson’s disease brains

We previously captured proteomic data^43^ from post-mortem Parkinson’s disease brain samples and we investigated alterations in the MAPK signalling pathway in human brain tissue (Supplementary Figure 4A). We found that mitogen-activated protein kinase 1 (MAPK1) was increased by 1.24-fold compared to healthy controls in Braak stage 3/4 substantia nigra, a region of the brain that is severely affected by disease at early stage, and 1.31-fold in early PD parahippocampus, a region that is only mildly affected at early stage (Supplementary Figure 4Bi&Bii). A related kinase, dual specificity mitogen-activated protein kinase kinase 1 (MAP2K1) also demonstrated upregulation in the early substantia nigra (1.80-fold) and the late frontal cortex (1.97-fold), a region that is only affected by pathology at later stages of the disease (Supplementary Figure 4Biii&iv).

Furthermore, when assessing the proteomic changes within the insulin receptor signalling pathway within the Ingenuity Pathway Analysis tool we observed multiple expression changes for proteins within the pathway in early PD substantia nigra compared to controls, leading to alterations in fatty acid synthesis, protein synthesis, apoptosis, sodium transport and transcription (Supplementary Figure 4C). These results together highlight that these pathways are also implicated within early stage sporadic PD post-mortem brain tissue and supports interventions that target these pathways.

### GLP1-R agonist in patients is associated with alterations in insulin signalling, protein aggregation and inflammation

Exenatide has previously been evaluated in PD in a 60-week, Phase II randomised control trial (NCT01971242) in which patients were treated with exenatide or placebo for 48 weeks, with collection of biological samples and clinical data every 12 weeks (Figure 6A). We previously reported exenatide treatment was associated with increased expression of neuronal IRS-1p Tyr, IRS-1 p-S616 and IRS-1 p-S312 in Neural-derived extracellular vesicles (NDEVs), and elevated expression of downstream substrates, including total Akt and phosphorylated mechanistic target of rapamycin (mTOR), which were also significantly associated with clinical improvement^44^.

**Figure 6.**
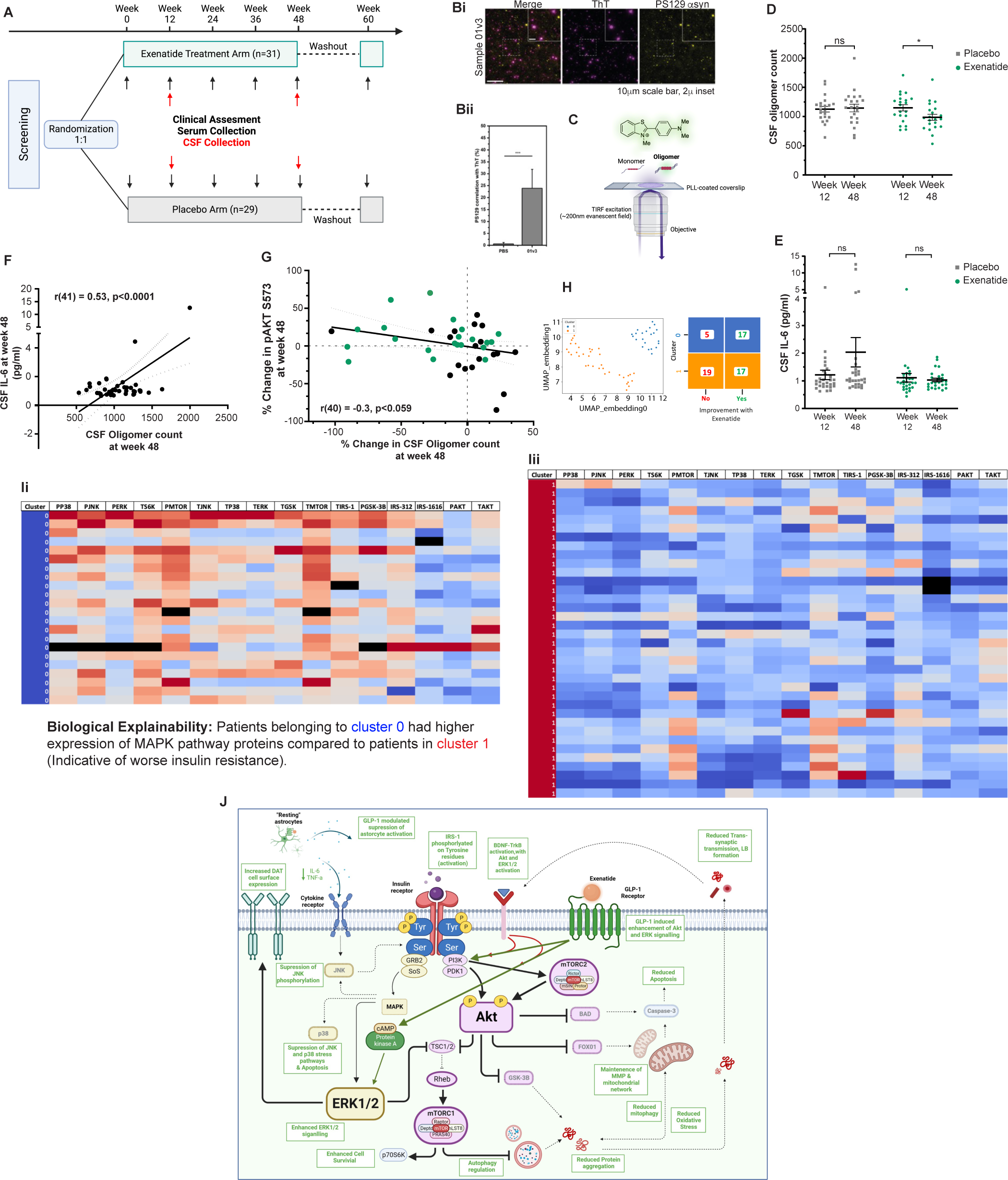
Analysis of biomarkers from the Exenatide-PD2 trial. **A.** Flow diagram showing study design of the Phase 2 RCT Exenatide-PD2 clinical trial. **Bi.** Representative image of single-molecule pull-down (SiMPull) detecting PS129 α-synuclein aggregates present in patient CSF. Magenta shows SAVE imaging with ThT, and yellow is **Bii.** Coincidence quantification of ThT and pS129 staining. **C.** Schematic of single-molecule visualization by enhancement (SAVE) imaging used for the detection of amyloid aggregates. **D.** Quantification of CSF oligomer count shows patients treated with exenatide had a significant reduction in CSF α- synuclein oligomer counts from Week 12 to Week 48 compared to controls. After adjusting for differences in gender, age, and baseline disease severity, there was a statistically significant interaction between the group (placebo/exenatide) and time on CSF Oligomer counts, F(1, 39) = 5.499, p =0.024, partial η2 = 0.126. While there was no significant difference between CSF oligomer counts at Week 12 (Placebo 1116.5 ± 54.4 vs Exenatide 1137.1 ± 55.8, p= 0.802), at Week 48, the CSF Oligomer count in the Placebo group remained unchanged (1159.5 ± 60. 5), while the Exenatide-treated group had a reduction in CSF Oligomers (949.4 ± 61.5), resulting in significant difference of – 210.0 (95%CI –392.2 - -27.8; p=0.025) **E.** IL-6 concentrations in the CSF were quantified at Week 12 and Week 48. There was no significant difference between CSF IL-6 concentration at Week 12 (Placebo 1.22pg/ml ± 0.1 vs Exenatide 1.11pg/ml ± 0.1, p= 0.689), but at Week 48, the CSF IL-6 concentration in the Placebo group increased to 2.08pg/ml ± 0.4, while the Exenatide-treated group had a reduction in CSF IL-6 (0.99 ± 0.4), resulting in a difference of –1.01pg/ml (95%CI –2.33 - 0.154; p=0.085) **F.** Pearson’s correlation between CSF oligomer counts and CSF IL-6 at Week 48 indicating a significant positive correlation between the CSF α-synuclein oligomer counts and CSF IL-6 r(41) = 0.53, p<0.0001. **G.** Pearson’s correlation between <u>CSF change in pAkt S573</u> and <u>change in CSF α-</u> <u>synuclein</u> oligomer counts at week 48 indicating a significant negative correlation r(40) = -0.30, p=0.059. **H.** Applying dimensionality reduction to the Neural-Derived Extra-cellular Vesicle biomarkers at baseline using UMAP identified 2 clusters of patients; Cluster 0, n=22, Cluster 1, n=37. **I.** Patients belonging to cluster 0 were predicted at baseline to show improvement with Exenatide compared to patients in cluster 1; Odds ratio 1.75, p=0.0280**. J.** Permutation testing to see which variables are significantly associated with the difference between cluster 0 and cluster 1 highlighted elevated levels of MAPK pathway kinases as predictors of response to exenatide. **J.** Diagram showing the restoration of neuronal insulin signalling by exenatide. Note: Statistical significance measured by Two-way ANCOVA with Bonferroni’s multiple comparisons test. *p < 0.05, ***p < 0.0005. Scale bar = 10µm.

#### Protein aggregation

Cerebrospinal fluid was collected from trial participants at week-12 and week-48 for the primary aim of detecting whether peripherally administered exenatide is detectable in the CSF. We utilized amyloid-binding dye thioflavin-T (ThT) and single-molecule visualization by enhancement (SAVE) imaging^45^ to quantify protein aggregates in the CSF at week-12 and week-48. SAVE imaging enables the detection of any amyloid aggregate, regardless of its protein composition. We therefore used a single-molecule pull-down (SiMPull) approach to specifically immobilize phosphorylated α-synuclein present in CSF, and probed this using labeled pS129-α-synuclein antibody (Figure 6Bi,Bii&C). We confirmed aggregates detected in CSF contained PS129 α-synuclein. A two-way mixed ANCOVA model was run to determine *if there were any significant differences in CSF oligomer concentration between the treatment armsover time.* While the CSF Oligomer count in the Placebo group remained unchanged, the Exenatide-treated group had a <u>significant reduction in CSF Oligomers</u>, resulting in significant difference between groups (Figure 6D)

#### Inflammation

IL-6 concentrations in the CSF were quantified at Week 12 and Week 48. There was no significant difference between CSF IL-6 concentration at Week 12, but at Week 48, the CSF IL-6 concentration in the Placebo group increased, while the Exenatide-treated group had a trend to reduction in CSF IL-6 (Figure 6E)

Furthermore, there was a significant positive correlation between CSF oligomer counts and CSF IL-6 at Week 48; (Figure 6F), suggesting those with higher CSF IL-6 also had a higher CSF aggregateload. Additionally, there was also a negative correlation between the % change in CSF aggregate load at week 48, and the % change in pAkt S573; (Figure 6G), indicating those patients with the greatest pAKt increase had the greatest reduction in CSF aggregate load.

### Higher MAPK kinase expression can predict a clinical response to exenatide

To explore predictors of exenatide response in trial participants, we defined Responders as participants who scored >2 points improvement over baseline in MDS-UPDRS III score at 48 weeks. Multimodal datasets from the Exenatide-PD2 trial including clinical outcomes and NDEV biomarker data were used to stratify participants to predict drug response at baseline based on these newly identified biomarker signatures (Supplementary Figure 5A for overview). The clustering analysis encompassed all participants at baseline, employing 5 PCA components that cumulatively accounted for over 90% of the variance. Subsequently, UMAP was applied for additional dimensionality reduction, followed by HDBSCAN for 2D UMAP clustering. This process revealed two distinct clusters (Figure 6H) – Cluster 0, comprising 22 participants, and cluster 1, comprising 36 participants. The HDBSCAN for 2D UMAP clustering was iterated 100 times with varied random initiations to ensure robustness of the clusters, consistently identifying the same participants in the same clusters 80% of the time (Supplementary Figure 5B). At baseline, individuals in Cluster 0 exhibited a 75% increase in the odds of being a “responder” to exenatide treatment compared to those in Cluster 1 (Figure 6I). Notably, among the 22 individuals in Cluster 0, all 17 of the “responders” (100%) were in the exenatide-treatment arm. Reassuringly, of the 5 non-responders in Cluster 0, 4 were in the placebo arm and only 1 individual was in the exenatide treatment arm (although this individual was later diagnosed with Progressive Supranuclear Palsy). To better understand the features driving the two clusters, we next employed permutation testing to identify variables significantly associated with the difference between Cluster 0 and Cluster 1. This analysis revealed that elevated levels of MAPK pathway kinases were key predictors of the two clusters.

Additionally, we conducted similar clustering models focusing solely on exenatide-treated participants, identifying two clusters that predicted treatment response at baseline. Critically, 100 iterations of HDBSCAN with varied random initiations resulted in the same two clusters on every occasion (Supplementary Figure 5D). Among those in Cluster 0, all participants (100%) showed improvement with exenatide treatment, compared to only 60% in Cluster 1 (Supplementary Figure 5E). Permutation testing on this model, found that similar elevated MAPK kinases were driving the two clusters (Supplementary Figure 5F). This is relevant as elevated levels of MAPK kinases have long been associated with insulin resistance, and supports our *in vitro* data, that exenatide treatment suppresses MAPK kinase activation. Although this requires further validation in clinical trials, individuals with marked neuronal insulin resistance at baseline may benefit the most from this class of drugs.

## Discussion

Insulin plays a vital role in neurodegeneration via binding to its receptor, which is found abundantly in the striatum, hippocampus and the cerebral cortex^46,47^. Insulin signalling is critical to neuronal survival in the brain, and maintains a balance between several pro-survival or pro-apoptotic pathways via downstream effectors such as mTOR, FOXO, NFkB, Caspase-3 and GSK-3B^8,48^. Thus, insulin resistance/dysfunctional insulin signalling can potentially lead to the dysregulation of a number of biological processes involved in the pathogenesis of PD, ultimately leading to cellular dysfunction and neurodegeneration (Figure 1A) ^49–51^. The actions of insulin in neurons are mediated through two major pathways – Akt and MAPK. Akt signalling acts as a master regulator of cell survival, metabolism, protein homeostasis, and is the central hub of both insulin and GLP-1 signalling. In PD, selective loss of dopaminergic neurons is accompanied by a marked decrease of pAkt S473 expression in the PD brain ^20,21,52^ and inhibition of the Akt pathway promotes neurodegeneration ^52–54^. The PI3K–AKT signalling pathway (via insulin) provides important inhibitory input to FOXO, which are transcription factors that coordinate adaptive responses to stimuli such as oxidative, metabolic and genotoxic stress. FOXO has been implicated in PD pathogenesis in GWAS studies ^55^; in the pathogenesis of “mitochondrial” PD ^56^ , and FOXO inhibition has protective effects on dopaminergic survival ^57,58^. Separate to the PI3K-AKT-FOXO pathway, insulin and GLP-1R stimulation also activates the MAPK pathway, which comprises three major branches, the JNKs, the p38 kinases and the ERKs. MAPK cascade signalling proteins can either modulate cell growth/proliferation, via activation of ERKs to direct cell growth and proliferation^21,59^, or MAPK may induce stress pathways via JNK and p38, which are activated by oxidative and cellular stresses to modulate inflammation and apoptosis. There is extensive crosstalk and feed-back loops between these branches, and between the Akt and MAPK arms.

We demonstrate evidence of neuronal insulin resistance and dysfunctional insulin signalling in iPSC-derived cortical and midbrain neurons derived from patients with *SNCA* mutations (i) *SNCA* A53T mutant and SNCAx3 midbrain dopaminergic neurons express significantly elevated IRS-1p S312 compared to control neurons, an indirect marker of insulin resistance^60,59^ (ii) Direct response to exogenously applied insulin results in a physiologically expected increase in Akt phosphorylation in control neurons, but significantly blunted downstream responses in *SNCA* neurons (akin to insulin resistance) (iii) Transcriptomic analysis revealed <u>three</u> (of the top six pathways) differentially expressed in *SNCA* midbrain dopaminergic neurons are involved in incretin (GLP-1/GIP) synthesis, secretion and inactivation and (iv) marked elevation of genes promoting/associated with insulin resistance, including elevated expression of p90 ribosomal S6 kinase (RPSKB1) – a key promoter of insulin resistance previously identified through GWAS and in vitro studies^61,62^. Consistent with evidence of “upstream” dysfunctional insulin signalling in *SNCA* mutant neurons, there is significantly reduced gene expression of pro-survival kinases Akt, RPTOR (regulatory associated protein of mTOR, complex 1), and BDNF, and elevated FOXO1 and Caspase-3 gene expression in these neurons. These transcriptional changes were associated with alterations in protein levels of these components of the Akt and MAPK pathways. Furthermore, we validated some of these findings in sporadic PD brain tissue, where we observed over-expression of MAP2K1, in the substantia nigra, which was identified as one of the top 3 genes overexpressed in *SNCA* mDA neurons. MAPK2K promotes oxidative stress and insulin resistance, suggesting this may play a role in PD pathogenesis of inherited and sporadic disease.

We have previously reported the temporal evolution of cellular abnormalities in *SNCAx3* and A53T mDA neurons^28,29,38–40^. Insulin resistance in *SNCA* mutant neurons was associated with mitochondrial fragmentation, elevated oxidative stress, perturbations in the autophagic and lysosomal pathway, and increased accumulation of aggregated α-synuclein, resulting in reduced cellular viability. Taken together, we hypothesise that the loss of cellular homeostasis, as a result of dysfunctional insulin signalling on the Akt and MAPK pathways, may, through increased FOXO activation, lead to oxidative stress, mitochondrial dysfunction, promotion of α-synuclein aggregation, activation of inflammatory pathways, downregulation of DAT receptors, and further promotion of IRS-1 phosphorylation on serine residues – perpetuating a vicious cycle of cellular stress and cell death (Figure 1A).

GLP-1 agonists, including exenatide, have demonstrated neuroprotective effects in *in-vivo* toxin models of PD ^22,63^, though a unifying mechanism remains uncertain. Here, given the inherent dysfunctional insulin signalling in *SNCA* mutant neurons, and the increasing evidence linking insulin resistance and neurodegeneration, we focused on its primary effects as an “insulin sensitising agent”. Treatment of *SNCA* mutant neurons with exenatide restored appropriate physiological responses to exogenous insulin and reduced IRS-1 s312 expression, thus providing further evidence that exenatide can restore inherently dysfunctional insulin pathways. Exenatide treatment was also accompanied by appropriate enhancement of “pro-survival” downstream signalling – with restoration/elevation of Akt phosphorylation, mTOR, and also ERK1/2 and BDNF expression, and suppression of “pro-apoptotic” FOXO1, Caspase-3, p38 and JNK expression (Figure 6J).

Our observations that exenatide restores both Akt and ERK1/2 signalling is novel, and in line with previous reports that suggest GLP-1R-induced activation of both Akt and ERK1/2 is necessary to confer beneficial effects on oxidative stress and mitochondrial function in *in-vitro* and *in-vivo* models^64^.

Exenatide treatment of *SNCA* neurons led to significant suppression of MAPK pathway kinases JNK and p38. JNK signalling is activated in response to inflammatory stimuli, and has been found to be elevated in postmortem PD brains ^65^, whereas pharmacological inhibition of JNK by GLP-1R agonists is protective in animal models of PD^59^, and a number of studies indicate degeneration of nigral dopaminergic neurons is accompanied by an increase in the level of p38 MAPKs ^67,68^, while pharmacological inhibition of p38MAPK prevents toxin-induced apoptosis in PD models ^69–72^.

Furthermore, overactivation of MAPK pathway kinases has long been associated with insulin resistance with a number of “insulin-sensitising drugs” acting to inhibit the MAPK pathway^73^

FOXO1 was highlighted as significantly overexpressed in mDA from patients with *SNCA* mutations, and is associated with exacerbating inflammation and apoptosis in PD pathogenesis. Exenatide acts to normalize FOXO1 phosphorylation (and reduce its activity) in *SNCA* neurons to control levels.

Exenatide-induced preservation of mitochondrial networks and normalization of mitophagy were associated with exenatide-induced increase in PI3K-Akt activation. This suggests that in dopaminergic neurons, beneficial effects on oxidative stress and mitochondrial function are probably associated with exenatide-enhancement of the PI3K-Akt-FOXO1 pathway.

Exenatide-induced increased expression of phosphorylated Akt S473 and ERK1/2, also modulates mTOR, as exenatide significantly elevates expression of mTOR in *SNCA* neurons. A key downstream kinase of Akt, mTORC1, is sensitive to insulin levels, and acts to promote protein synthesis and supress autophagy ^76,77^, while inactivation of mTOR by reduced Akt activation, triggers the autophagy cascade. Supporting this are multiple studies which have shown that pharmacological suppression of mTOR by neurotoxins enhances neurodegeneration^78^. Studies utilising neuronal cultures and mice expressing mutant *A53T* α-synuclein indicate mTORC1 signalling is overactivated, resulting in insulin resistance (via IRS-1 serine phosphorylation)^79^ and that GLP-1 agonists can restore dysfunctional autophagy ^63^, via the PI3K/Akt/mTOR pathway ^80–82^. Thus, exenatide may positively influence the balance between IRS-1 inhibition (mTORC1 directed negative feedback) and Akt responsiveness to insulin (mTORC2 directed positive feedback) to favour restoration of homeostatic insulin signalling.

Taken together, exenatide-induced restoration of intracellular signalling was associated with positive observed biological effects in mDA neurons including preservation of mitochondrial networks, reduction in oxidative stress, normalisation of lysosomal activity, and reduction in aggregation of α- synuclein in *SNCA* neurons, ultimately resulting in preservation of cellular viability (Figure 6J).

We have established neuron-specific effects of insulin resistance, and aberrant downstream pathways. However, GLP-1R agonism has been previously shown to have effects on glia^83^. α- synuclein can induce pathogenic activation of midbrain astrocytes, characterised by inflammatory transcriptional responses, downregulation of phagocytic function, via activation of TLR4 on the cell membrane and downstream activation of IKK-NF-κB and MAPK signalling cascades^84–86^. We show astrocytes derived from patients with *SNCA* mutations express significantly greater levels of inflammatory cytokines IL-2 and IL-6 at baseline compared to control astrocytes typical of an inflammatory state, and produce a significantly greater inflammatory response when exposed to α- synuclein oligomers or LPS compared to control astrocytes; and are “primed” to be reactive.

However, pre-treatment of *SNCA* astrocytes with exenatide reverted the “inflammatory A1” profile of basal *SNCA* astrocytes, to a less reactive phenotype, characterised by significantly reduced inflammatory cytokines IFN-y, IL-6 and TNF-a, with a subsequent prevention of neuronal cell death. These protective effects were abolished in the presence of Ex9-39 (a GLP-1R antagonist) – suggesting the GLP-1R mediates these effects. Pre-clinical studies from MPTP, 6-OHDA and α-synuclein PFF rodent models of PD, demonstrate potent anti-inflammatory effects of GLP-1R agonists that may be due to direct interaction with the GLP-1R present on microglia and astrocytes, leading to suppression of microgliosis and astrogliosis, shifting glial cells to a “protective” state, with associated inhibition of TLR4, Iba-1, GFAP, NF-κB, TNF-α and IL-6 ^87–90^. GLP-1 itself is both a target and mediator of inflammation^91–94^, and GLP-1R expression is upregulated in response to inflammatory states ^95,96^, hence offering the advantage of avoiding the potentially deleterious consequences of other “anti-inflammatory” treatments that broadly suppress physiological responses of microglia and astrocytes to injury.

We previously evaluated exenatide for the treatment of PD in a clinical trial – Exenatide-PD2 (NCT01971242), which demonstrated participants treated with exenatide had motor and non-motor benefits compared to untreated participants^97^. We utilized CSF and neuronal-derived extracellular vesicles (NDEVs) from participants’ serum as source of biomarkers to identify any evidence of target engagement/drug response, that aligned in both patients and our human *SNCA* neuronal models. Participantstreated with exenatide also had significantly elevated levels of neuronal insulin signalling proteins (IRS-1 pTYr, IRS-1 s312, IRS-1 s616), as well as elevated pAkt and mTOR^98^. Clinical motor improvements observed in the exenatide treated group were at least partially explicable by concomitant elevation of mTOR and pAkt expression. We also observed similar increases in pAkt signalling and mTOR in *SNCA* iPSC-derived neurons following exenatide treatment. Finally, in order to support our *in-vitro* observations that exenatide reduces accumulation of α-synuclein, we deployed a highly sensitive single-molecule assay that can detect individual small aggregates, or oligomers, of α-synuclein. At week 48 of the trial, exenatide treated patients had a significant reduction in CSF oligomers compared to the placebo group. In addition, there was a significant negative correlation observed between pAkt (derived from participant NDEVs) and CSF oligomer load. That is, as (exenatide-induced) pAkt increases, CSF oligomer load decreases, suggesting that the elevation / restoration of pAkt by exenatide is ultimately associated with reduced aggregation of α-synuclein. Supporting the *in-vitro* anti-inflammatory effects of exenatide treatment, there was a trend towards lower CSF IL-6 in the exenatide-treated group compared to the placebo group.

Taken together, this suggests that through its restoration of insulin signalling, exenatide influences the aggregation of α synuclein, shown here in our *in-vitro* models as being associated with increased cell survival, and in participants from the Exenatide-PD2 trial, with associated clinical benefits. It further highlights the utility of developing biomarkers for trials that reflect the pathogenesis of disease and therefore help distinguish ‘disease modifying’ therapy from purely symptomatic ones.

In order to predict patients that were potential responders to exenatide, we conducted an exploratory unsupervised machine learning analysis of clinical response and biomarker data from the Exenatide-PD2 trial. We were able to identify (at baseline prior to trial commencement), a cluster of 17 individuals (of all 60 participants) and a cluster of 11 individuals (of 29 exenatide treated participants) that were accurately predicted to show significant improvement with exenatide.

Elevated levels of MAPK pathway kinases (associated with insulin resistance, oxidative stress) at baseline were driving these clusters. Our in vitro platform demonstrates that exenatide acts to supress MAPK pathway kinases, and therefore we hypothesis that participants with worse “neuronal insulin resistance”, and elevated levels of MAPK pathway kinases at baseline, are more likely to improve / respond to GLP-1R agonists that aim to restore dysfunctional insulin signalling / brain metabolism. Using NDEVs to measure neuronal insulin resistance could be potentially utilised to stratify and target individuals for disease modifying trials involving these classes of drugs in the future.

## Conclusions

We have previously observed exenatide treatment in a Phase 2 clinical trial of Parkinson’s led to motor and non-motor benefits, which were associated with modulation of insulin signalling and downstream Akt pathway kinases. Here, we demonstrate exenatide restores inherently dysfunctional insulin signalling in *SNCA* neurons with subsequent activation of the PI3K-Akt and suppression of MAPK-ERK pathways via a GLP-1R-dependent mechanism. Exenatide exerts multiple pleiotropic effects on neuronal function, through effects on both neurons and on glia, ultimately leading to reduction in aggregates of α-synuclein and promotion of cell survival. Similar pleiotropic effects were observed in biological samples from participants of the Exenatide-PD2 clinical trial, where exenatide led to a significant reduction in α-synuclein aggregatesin the CSF, and a reduction in inflammatory cytokine IL-6, mirroring our observations in iPSC models. These data provide a mechanism by which GLP-1 agonism can alter underlying disease pathology in PD, and further support therapeutic interventions aimed at restoring insulin sensitivity in the treatment of Parkinson’s.

## Acknowledgements

We would wish to thank the patients for the fibroblast donation and for their participation during the Exenatide-PD clinical trials. We would also like to thank the Francis Crick Institute Flow Cytometry, Advanced Sequencing, and Bioinformatics and Biostatistics STPs for their help in conducting and analysing the flow cytometry, and single-cell RNA-seq experiments. We also wish to thank Cure Parkinson’s Trust for their help and support during the Exenatide-PD clinical trials. This research was funded in whole or in part by Aligning Science Across Parkinson’s (grant ASAP-000509) through the Michael J. Fox Foundation for Parkinson’s Research (MJFF). For the purpose of open-access, the author has applied a CC public copyright license to all Author Accepted Manuscripts arising from this submission.

## Funding

DA acknowledges funding from the Cure Parkinson’s Trust and NIHR. GV acknowledges funding from the UCL-Birkbeck MRC DTP. KI is part funded by National Institute for Health and Care Research (NIHR) Maudsley Biomedical Research Centre (BRC). Dr Davies has acted as a consultant, advisory board member, and speaker for Boehringer Ingelheim, Eli Lilly, Novo Nordisk, and Sanofi; has served as an advisory board member and speaker for AstraZeneca; has served as an advisory board member for Medtronic, Pfizer, and ShouTi Pharma; has served as a speaker for Amgen, Novartis, and Sanofi; and has received grants in support of investigator and investigator-initiated trials from AstraZeneca, Boehringer Ingelheim, Eli Lilly, Janssen, Novo Nordisk, and Sanofi. TF acknowledges funding from Cure Parkinson’s, National Institute for Health Research, MJFox Foundation, John Black Charitable Foundation, Janet Owens Research fellowship, Edmond J Safra Foundation, Van Andel Institute, and Defeat MSA. SG acknowledges funding from the i2i grant (The Francis Crick Institute), MJFox foundation, Wellcome. SG is an MRC Senior Clinician Scientist. This work is supported by the Francis Crick Institute which receives funding from the UK Medical Research Council, Cancer Research UK, and the Wellcome Trust. RS, MHH and CL acknowledge funding from UCB Biopharma, an EPSRC IAA award. RS is an MND Association Lady Edith Wolfson Junior Non-Clinical Fellow (Saleeb/Oct22/980-799 (RSS)). JO’S and MHH acknowledge funding from the National Institutes of Health (5-r01-NS127186-02). The TIRF microscope used in this study was funded by UCB Biopharma, the UK Dementia Research Institute, and a kind donation from Dr. Jim Love. CL was funded by a British Heart Foundation Research Excellence Award. The Exenatide PD2 trial was hosted by the Leonard Wolfson Experimental Neuroscience Centre, Queen Square Institute of Neurology

## Data availability

The data that support the findings of this study are available on Zenodo (upload in progress). Single-cell RNA-seq raw data were deposited to EGA. The code used for all scRNA-seq and RNA velocity analysis has been deposited online. The CellProfiler pipeline can be downloaded here: https://cellprofiler.org/published-pipelines (available on publication)

## Materials and Methods

### Human induced pluripotent stem cell culture

hiPSC culture hiPSCs were derived from donors who had given signed informed consent for the derivation of hiPSC lines from skin biopsies as part of the EU IMI-funded program StemBANCC and reprogrammed as described (Devine et al., 2011; Little et al., 2018). Briefly, the CytoTune-iPS reprogramming kit (ThermoFisherScientific) was used to reprogram fibroblasts through expression of OCT4, SOX2, KLF4 and c-MYC by four separate Sendai viral vectors. Control 1 and 2 were derived by StemBANCC from an unaffected volunteer, and control 3 was purchased from ThermoFisherScientific. hiPSCs were generated from familial PD patients carrying a point mutation in *SNCA* (*A53T* point mutation), and from a patient with early onset autosomal dominant PD owing to a triplication of the *SNCA* gene (*SNCA*x3). The presence of mutations was confirmed by sanger sequencing (GENEWIZ). hiPSCs were maintained on Geltrex in Essential 8 medium or mTeSR (ThermoFisherScientific) and passaged using 0.5 mM EDTA. All lines were mycoplasma tested (all negative) and performed with short tandem repeat profiling (all matched) by the Francis Crick Institute Cell service team. The Isogenic control lines of *SNCA A53T* and *SNCA*x3 were generated using CRISPR/Cas9 editing by Applied StemCell Inc. (USA, project ID: C1729).

### Differentiation into cortical neurons

Neural induction and differentiation were performed using a modified, published protocol (Shi et al., 2012). Briefly, upon 100% confluence of hiPSC, dual SMAD inhibitors SB431542 (10 µM, 997 Tocris) and Dorsomorphin (1 µM, Tocris), were added to N2B27 medium (DMEM:F12, insulin, 2-mercaptoethanol, Non Essential Amino Acids, N2 supplement, Pen/Strep, Neurobasal, B27, Glutamax, Pen/Strep) and the medium was refreshed every day for 10-12 days until neuroepithelium appear. Then the neuroepithelium sheets were split using dispase and cultured in N2B27. At around 35 days of induction, cells were dissociated into a single cell using accutase and approximately 150,000 number of cells plated on either PDL and laminin coated glass bottom 8-well slide chambers (Ibidi/Thistle, cat No.80826), Geltrex-coated 8-well ibidi chambers (cat No. IB-80826) or 96-well plates (Falcon, cat No. 353219). The medium was replaced every 4-5 days 34 and cells were used at 60-90 days after induction. Neuronal cultures were verified using immunocytochemistry for neuronal markers, MAP2 and TRB1 (see the antibody lists in Immunohistochemistry). Expression of glutamate receptors and voltage-gated calcium channels was confirmed by measuring the calcium response to physiological glutamate concentrations.

### Differentiation into midbrain dopaminergic neurons

Differentiation was triggered by removing old media and replacing it with a chemically defined medium consisting of DMEM/F12, N2 supplement, Neurobasal, B27 supplement, L-Glutamine, non-essential amino acids, 50U/ml penicillin-streptomycin, β-mercaptoethanol (all from ThermoFisherScientific) and 5μg/ml insulin (Sigma), termed “N2B27”. Cells were patterned for 14 days with daily media changes. For the first 2 days, the media was supplemented with the small molecules 5μM SB431542 (Tocris Bioscience), 2μM Dorsomorphin (Tocris Bioscience), 1μM CHIR99021 (Miltenyi Biotec). On day 2, 1μM Purmorphamine (Merck Millipore) was added. On day 8, CHIR99021, and SB431542 were removed leaving only Dorsomorphin and Purmorphamine in the medium until day 14. Cells were enzymatically dissociated and split on days 4, 10, and 14 using 1mg/ml of Dispase (ThermoFisherScientific). After patterning, mDA neuronal precursor cells (NPCs) were maintained in N2B27 for 4 days. On day 19, cells were plated onto Geltrex pre-coated Ibidi 8-well chambers (100k/well), clear bottom 96-well plates (50k/well), or 12-well plates (500k/well), and terminally differentiated using N2B27 supplemented with 0.1μM Compound E (Enzo Life Sciences) and 10μM Y-27632 dihydrochloride (Rho kinase ROCK inhibitor) (Tocris) from day 20 for the whole duration of terminal differentiation, with two weekly media changes.

### Differentiation of hiPSC into astrocytes

Differentiation of cortical region-specific astrocytes was performed using a modified protocol^30,31^. Briefly, as demonstrated in Fig 1b, hiPSC were differentiated into neural precursor cells (NPCs) using an established protocol (Shi et al., 2012). To derive glial precursor cells (GPCs), NPCs were cultured with dual SMAD inhibition for 25-30 days, followed by culturing with the neural induction medium supplemented with 20 ng/ml human FGF-2. The passage was performed twice per week (1:2 or 1:3) using Accutase (Cat #A1110501, Thermo Fisher Scientific) by vigorously breaking pellets to remove neuronal cells. Upon the appearance of glial morphology (around day 90 from the neural induction), the GPCs were cultured for 7 days with 10 ng/ml bone morphogenetic protein 4 (BMP4) and 20 ng/ml leukemia inhibitory factor (LIF) which activates the JAK/STAT signalling pathway, refreshing the medium every other day. On the 8th day, BMP4 and LIF were withdrawn, and the GPC were further differentiated for maturation in the neural induction medium without human FGF-2. 3-4 times more passages are required (1:3 or 1:4) until the complete loss of the precursor property, including proliferation.

### Collecting cell pellets for bulk RNA-seq

To collect cell pellets, samples were trypsinised or scraped from the culture surface and placed in a 15ml conical tube. These tubes were centrifuged at 800g in a refrigerated centrifuge for 5 minutes, and the culture media decanted. The pellet was resuspended in 10ml chilled PBS per tube by pipetting, then centrifuged again using the above parameters before decanting the PBS. For bulk RNA sequencing, the cell pellets were frozen on dry ice and stored at -80°C.

### Collecting cell pellets for single cell RNA-seq

Samples for single cell RNA sequencing followed the procedure above, though instead of freezing, the pellets were resuspended in 1ml PBS, and 100,000 cells were transferred to a 1.5ml falcon tube. These were centrifuged at 1000 rpm for 3 min at 4C, before resuspending cells in 20ul chilled DPS. 180ul chilled 100% methanol was added dropwise to the cells while gently vortexing to prevent the

### Bulk Single-cell RNA-sequencing data generation and processing

Libraries for sequencing were prepared using the polyA KAPA mRNA Hyper Prep kit by loading 50 ng of total RNA into the initial reaction; fragmentation and PCR steps were undertaken as per the manufacturer’s instructions. Final library concentrations were determined using Qubit 2.0 fluorometer and pooled to a normalized input library. Pools were sequenced using the Illumina HiSeq Sequencing system to generate 75 bp single-end reads with an average read depth of ∼20million paired-end reads per sample.

### Single-cell RNA-sequencing data generation and processing

The reads were aligned to the human reference genome (Ensembl release 93, GRCh38) using Cell Ranger v3.0.2. The analysis was carried out using Seurat v3.0 (REF-Butler, 2018, Stuart, 2019 #645) following Seurat’s standard workflow. Cells expressing fewer than 200 genes were excluded from the subsequent analysis. In addition, we excluded cells with more than 3000 detected genes to remove suspected cell doublets or multiplets. Given that certain cell types, e.g. neurons, naturally express higher levels of mitochondrial genes, we applied a 10% cut -off for the percentage of mitochondrial genes expressed to filter out likely apoptotic cells. Using default parameters of Seurat, data for each sample were log normalised across cells and the 2000 most highly variable genes identified. Using the canonical correlation analysis (‘CCA’) [https://www.cell.com/cell/fulltext/S0092-8674(19)30559-8] to identify anchors, we integrated the samples using Seurat v 3 , followed by regression of the effect of cell cycle and scaling of the data. Dimensional reduction was performed using 50 PCs. We used Clustree v0.4.4 {Zappia, 2018 #646} and Seurat’s plots to visualise the expression of astrocytic and neuronal marker genes across different cluster resolutions (0.05 - 0.5 in 0.05 increments and 0.5 - 1.0 in 0.1 increments). A clustering resolution 0.25 was selected, as it was the lowest resolution with stable clusters of cells, that explained the heterogeneity in the samples. The differentially expressed genes between the clusters of interest were identified using Seurat’s FindMarkers() and the default ’Wilcox’ test.

### RNA Sequencing Analysis

Raw reads were processed using the RNA-seq nfcore pipeline (version 3.2) (Patel, 2021). The pipeline uses the nf-core framework with nextflow (version 21.04.0). In short, reads were trimmed using trimgalore (version 0.6.6) and subsequently aligned and quantified with STAR-RSEM (version 1.3.1) against the human genome GRCh38 and annotation release 95, both from Ensembl.

Differential gene expression analysis was performed in R-4.1.2 (R Core Team, 2021) using the DESeq2 package (version 1.34.0). Using the model ∼Neuronal_Induction + Cell_Genotype + Age_of_cells + Genotype the gene expression between SNCA and control genotypes was compared for each of the investigated timepoints during the differentiation into midbrain dopaminergic neurons. The regularised (“shrunken”) log2 fold changes for genes of interest were visualised as a heatmap using the ComplexHeatmap package (version 2.10.0).

Reactome pathway enrichment analysis was performed in R-4.1.2 using the ReactomePA package (version 1.38.0) with default parameters and using the visualisation functions from the clusterProfiler package (version 4.2.2).

### Immunohistochemistry

Cells were fixed in 4 % paraformaldehyde and permeabilized with 0.2 % Triton-X 100. 5 % BSA 1012 was used to block non-specific binding before cells were incubated with primary antibodies either for 2 hours at room temperature or overnight at 4 °C. The next day, cells were washed three times with PBS and incubated with secondary antibody for 1hr at room temperature. Cells were mounted with antifading medium after three times wash steps (DAPI was added in the second wash if required) and let dry overnight. 1017 Lists of primary antibodies used; Anti-MAP2 (abcam, ab183830, 1:500), Anti-TRB1 (abcam, 1018 ab137717), Recombinant Anti-α-synuclein antibody [MJFR1] (abcam, ab138501, 1:250), Recombinant Anti-α-synuclein aggregate antibody [MJFR-14-6-4-2] - Conformation-Specific (abcam, ab209538, 1:200). Lists of secondary antibodies used; Goat Anti-Chicken IgY H&L (Alexa Fluor® 488) (abcam, ab150169, 1:500), Goat Anti-Mouse IgG H&L (Alexa Fluor® 555) (abcam, ab150114, 1:500), Goat Anti-Rabbit IgG H&L (Alexa Fluor® 647) (abcam, ab150079, 1:500).

### Live-cell imaging

The samples were imaged using Zeiss 880 confocal system with a 40x, 1.4 N.A. oil objective, and a pinhole of 1 airy units (AU). Between 3-5 images were collected per sample, all with a Z projection consisting of >6 slices, and displayed as a maximum projection. Samples were also imaged using the PerkinElmer Opera PhenixTM High Content Screening System with 20 and 40x water objective lenses. A minimum of 3 fields of view and a Z projection of 6 slices was acquired per well, with the images displayed as a maximum projection. The accompanying software, ColumbusTM was used to store and analyse acquired images. The settings for the acquisition of images were kept the same for all samples in the experiment set. The data were analysed using Andor iQ software (Belfast, UK). For confocal microscopes, illumination intensity was limited to 0.1-0.2% of laser output to prevent phototoxicity, and the pinhole was set to allow optical slice at approximately 1-2 μm. Room temperature (RT) HBSS was used as a recording buffer. 3-4 fields of view per well and at least 3 wells per group were used to analyze using ZEISS ZEN software, Volocity 6.3 cellular imaging ImageJ or Cell Profiler software. Image analysis and quantification were performed using the open-source software CellProfiler. A CellProfiler pipeline was designed to detect and count live and dead cells, mitochondria and lysosomes from immunofluorescence images using automatic thresholding and segmentation methods (see All experiments were repeated at least 2-3 times with different inductions (hiPSC derived neurons).

### Measurement of Oxidative stress

Superoxide prodcution: cells were washed and loaded 2 uM dihydroethidium (HEt; Thermo Fisher Scientific) in the recording buffer. HEt is an indicator of superoxide which exhibits blue fluorescence in the cytosol before oxidation, and the nucleus presents a red fluorescence upon oxidation. Het allows the rate of superoxide generation to be measured which is present as the ratio of the oxidized form of the dye over the reduced form. The recording was performed using an epi-fluorescence inverted microscope equipped with 20x objective after a quick loading (2-3 min) in order to limit the intracellular accumulation of oxidized product, and the dye was present throughout the imaging. Excitation was at530 nm, and emission recorded above 560 nm was assigned to the oxidized form, while excitation at 380 nm and emission collected from 405 nm to 470 nm was assigned to the reduced form. The ratio of the fluorescence intensity, resulting from its oxidized/reduced forms, was quantified and the rate of ROS production was determined by dividing the gradient of the HEt ratio after application of recombinant α-Syn against the basal gradient.

### Multiplex assay (cell death, lysosomal dynamics, mitochondrial membrane potentia)

Cell death was detected using Propidium iodide (PI, Thermo Fisher Scientific) or SYTOX™ Green (SYTOX, Thermo Fisher Scientific) which is excluded from viable cells but exhibits red fluorescence following a loss of membrane integrity and Hoechst 33342 (Hoechst, Thermo Fisher Scientific) which stains chromatin blue in all cells to count the total number of cells. 20 μM PI or 500 nM SYTOX and 10uM Hoechst were directly added into the dishes and cells were incubated for 15 min. The fluorescence measurements were detected using confocal microscopy. Hoechst and PI 38 were excited by 405 nm and 543 nm laser lines with the emission between 405 nm to 470 nm and 570 nm to 640nmrespectively. SYTOX was excited by a 488 nm laser with emission between 488 nm and 516 nm. The percentage of cell death was quantified by the number of red (PI) or green (SYTOX) fluorescent cells 33342 expressing cells per image. This data reflects the proportion of cells measured, and the ratio is compared across experimental conditions. To measure mitochondrial membrane potential, cells were incubated with 25nM tetramethylrhodamine methyl ester (TMRM, ThermoFisherScientific) in HBSS for 40 minutes and then imaging was acquired using Zeiss LSM 880 confocal microscope. TMRM is a lipophilic cationic dye that allows visualisation of mitochondria and accumulates within mitochondria in inverse proportion to mitochondrial membrane potential (Δψ_m_). The 560 nm laser line was used to excite and it was measured above 560 nm. Approximately 3-5 fields of view with Z projections were taken per sample. To measure lysosomal dynamics, cells were incubated with 50nM LysoTrackerTM Deep Red (ThermoFisherScientific), and Hoechst 33342 in HBSS for 40 minutes and then imaged using Zeiss LSM 880 confocal microscope where the 405 nm, and 647 nm laser line were used to excite Hoechst 33342 and LysoTrackerTM Deep Red, respectively. Approximately 4-5 fields of view with Z projections were taken per sample.

### Protein extraction

Cells were washed with cold PBS and then lysed with cold RIPA buffer (Thermo Fisher Scientific) supplemented with a cocktail of phosphatase and protease inhibitors (Life Technologies). After centrifugation, the supernatant was collected, and protein concentration was measured using a BCA or a Bradford quantification assay (Thermo Fisher Scientific). Equal amounts of protein lysates were used to perform ELISAs according to manufacturer’s instructions.

### Immunoassays

We used Meso Scale Diagnostics electrochemiluminescence assays to quantify IRS-1 pSer312, pAkt (Ser473), total Akt, p38, j-JNK and ERK1/2. 20µg of protein was loaded into each well in duplicate, and mean absorbance values were normalised to control values. Rigorous quality control assessments and procedures were used to ensure data quality. IRS-1 pSer312 was measured using the Phospho-IRS-1 (Ser312) Whole Cell Lysate Kit (with SULFO-TAG Anti-phospho (S312) antibody); pAkt and total Akt were measured using the Phospho(Ser473)/Total Akt Whole Cell Lysate Kit; phospho-JNK (Thr183/Tyr185), phospho-p38 (Thr180/Tyr182), and phospho-ERK-1/2 were measured using MAP Kinase Whole Cell Lysate Kit (with SULFO-TAG anti-total p38, anti total ERK1/2 and anti phosphor-JNK antibodies). All assays were used according to manufacturer’s instructions. All samples were run in duplicate, and coefficients of variance (CV) were determined for each sample. The mean CVs for the plate control for all analytes were under 20%. The limit of detection (LOD) was determined for each analyte by calculating the mean signal of the blank plus 2.5 times the standard deviation of the blank. Similarly, the lower limit of quantification (LLOQ) for each analyte was calculated using the mean signal of the blank plus 9 times the standard deviation of the blank; and recovery of the nearest standard between 80-120%. For IRS-1 pSer312, given that no standards were provided by the manufacturer of the assays, quality control had to rely exclusively on CVs. Therefore, for these markers, we adopted a more stringent threshold and all samples with electrochemiluminescence signal CV ≥ 15% were excluded from the analysis

### Modelling insulin resistance in vitro

Cells were cultured overnight using media supplemented with B27 without insulin (Thermofisher A1895601). After the overnight culture, insulin (Sigma Cat. 91077C) or vehicle was added to the media as indicated. Insulin was prepared fresh from frozen stocks every day before administration.

### SAVE imaging of protein aggregates

SAVE imaging was performed as previously described^45^. Borosilicate glass cover slips (22 x 40 mm, VWR, 6310135) were cleaned using an argon plasma cleaner (Zepto, Diener, Germany) for 1h to remove any fluorescent residues. Frame-Seal slide chambers (9 x 9 mm^2^, Biorad, Hercules, USA, SLF-0601) were affixed to the glass and 50 µL of poly-L-lysine (70k-150k molecular weight, Sigma-Aldrich, 25988-63-0) were added to the inside of the chamber on the glass coverslip and incubated for at least 30 minutes, before washing with 0.02 µm filtered PBS. CSF samples were diluted 10-fold in 0.02 µm filtered PBS and added to the glass cover slip (VWR, 6310135) and incubated for 1 hour. The sample was removed and the surface washed three times with filtered PBS, before 5 µM ThT (Abcam, ab120751) was added. Imaging was carried out on an ONI Nanoimager, using a 405 nm laser at 40% power and a TIR angle of 53.5°. Images were collected at 20 frames s^-1^ for a total of 50 frames. To reduce region bias, 25 images were collected spaced 200 µm apart, for three separate regions of the slide.

The data were analysed using custom-written code in Python 3.8 (code available at 10.5281/zenodo.7546532). The images were first averaged over the 20 frames, and the background subtracted using threshold_local in skimage.filters. Spots corresponding to ThT-bound aggregates were selected by applying a threshold equal to the mean + 5 x S.D. of the intensity in each image. The total number of aggregates were counted in each image and the data combined to give an overall count normalized to the imaging area.

### SiMPull imaging of protein aggregates

Cover slip passivation was carried out based on published procedures^99,100^. Briefly, plasma cleaned cover slips were incubated in 1M potassium hydroxide for 20 min, rinsed in deionised water and transferred to 1% (3-aminopropyl)trimethoxysilane diluted in methanol supplemented with 5% acetic acid for 20 min incubation. Following serial washing in methanol and deionised water, cover slips were dried with argon gas and attached to an 18-well gasket (Ibidi 81818). A solution of 100 mg/mL mPEG-SVA (5000 Da, Laysan Bio), 1 mg/mL biotinylated mPEG-SVA (5000 Da, Laysan Bio) and 0.1M NaHCO_3_ was applied for 16 hour incubation, then rinsed with deionised water and the glass dried with argon gas, before incubating 10 min with 0.2 mg/mL streptavidin in 0.02 um-filtered T50 buffer (10 mM tris pH 8.0 supplemented with 50 mM NaCl). 100 nM PS129 antibody (Abcam, ab209422) biotinylated in-house (Pierce, 90407) was conjugated to the surface via 20 min incubation. CSF was applied neat and incubated 24 hours, wells were washed with PBS and a 2 nM mix of PS129 conjugated to Alexa Fluor 488 and Alexa Fluor 647 (ThermoFisher, A20181/6) applied for 3 hours, rinsed and imaged on an ONI Nanoimager using a TIR angle of 53.5°. An 8 x 8 grid of 200 µm-spaced fields were imaged using sequential 638 nm and 488 nm excitation. The same regions were imaged with 405 nm excitation following application of 10 µM ThT (Abcam, ab120751), using the AF647 channel for image registration. Images were collected at 20 frames s^-1^ for a total of 20 frames in each channel.

The data were analyzed using custom-written macros in Fiji. Ground-truth Tetraspeck bead (ThermoFisher Scientific, T7279) image data was used to auto-align split channel images with the imreg python library and misalignments in stage position return pre- and post-ThT application were corrected by landmark registration in the AF647 channel. ThT and AF488 image series were maximum intensity projected and the ComDet plugin (v.0.5.5) for Fiji used for particle detection and the computation of event coincidence.

### Statistical analysis

Origin 2020 (Microcal Software Inc., Northampton, MA) software was employed for the statistical analysis and exponential curve fitting. An assessment of the normality of data was performed using Shapiro-Wilk and Kolmogorov Smirnov Normality tests. When the decision was made not to reject normality at 5% level, statistical tests were performed using unpaired two-sample t-tests, one-way ANOVA or two-way ANOVA corrected with a Bonferroni for multiple comparisons. Experimental data are shown as means ± standard error of the mean (SEM) or standard deviation (SD), and P value is set at 0.05. n = fields of view, if not stated otherwise. Sample sizes for experiments were selected to capture (1) technical variation including numbers of cell/field of view and coverslips, and (2) biological variations including numbers of the independent inductions and clones or patient line for hiPSC derived neurons. F-statistics was used to estimate variance within each group. Data were collected and analysed without bias. For live imaging experiments in human neurons, application of standard controls was used to determine the dynamic range of the signal measured (the maximum and minimum signal). The data are normalized to the maximum and minimum, and the same experimental conditions used to compare two states i.e. control vs *A53T* mutant. In the absence of a known standard in these experiments, comparisons across different conditions/cell lines are only relative to each other within this experimental paradigm.

## Notes

https://zenodo.org/records/10623671

### Competing Interest Statement

The authors have declared no competing interest.

