## Supplemental Material for "GLP1 receptor agonism ameliorates Parkinson’s disease through modulation of neuronal insulin signalling and glial suppression"

### Supplementary Materials

#### Supplementary figure 1 – GLP-1R agonist rescues neuronal insulin resistance in SNCA A53T

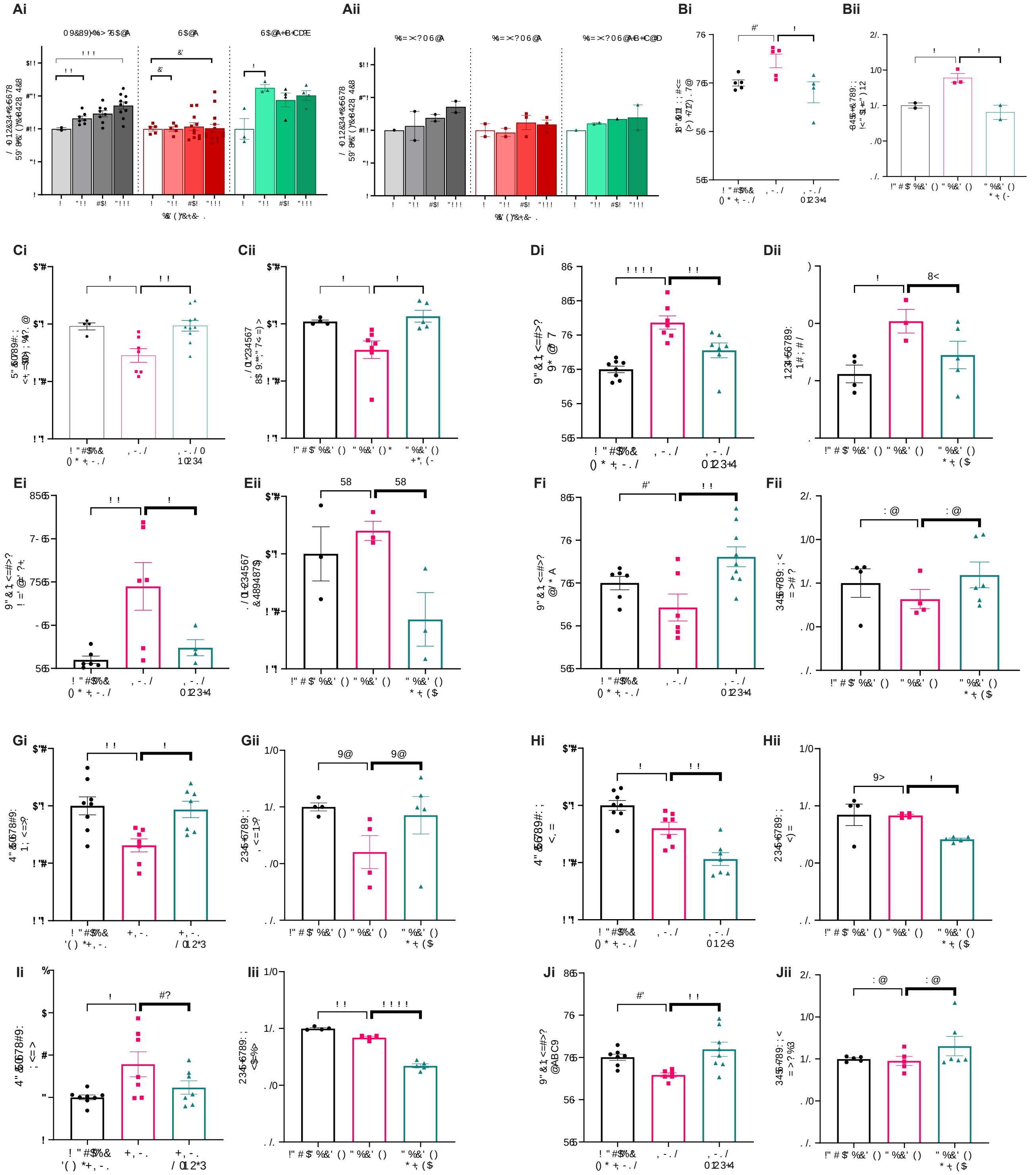


#### midbrain dopaminergic neurons (data split by cell line)

Supplementary figure 1

**A.** % changes in phosphorylated Akt S473 in control, A53T SNCA (**i**) and SNCA x3 (**ii**) mDA neurons in response to chronic insulin treatment +- Ex-4. **B.** Fold change of IRS-1 pS312 in A53T (**i**) and SNCAx3 (**ii**) mDA neurons. Fold changes quantification of kinases in the insulin signalling pathway in A53T (**i**) and SNCAx3 (**ii**) mDA:  **C.** p-AKT (S347), **D.** FOXO1, **E.** Caspase-3, **F.** mTOR. Fold change quantification of MAPK signalling pathway markers in A53T (**i**) and SNCAx3 (**ii**) mDA neurons: **G.** ERK1/2, **H.** p38, **I.** p-JNK, **J.** mBDNF.

### Supplementary figure 2 - Reactome pathways differentially expressed between SNCA and control mDA neurons

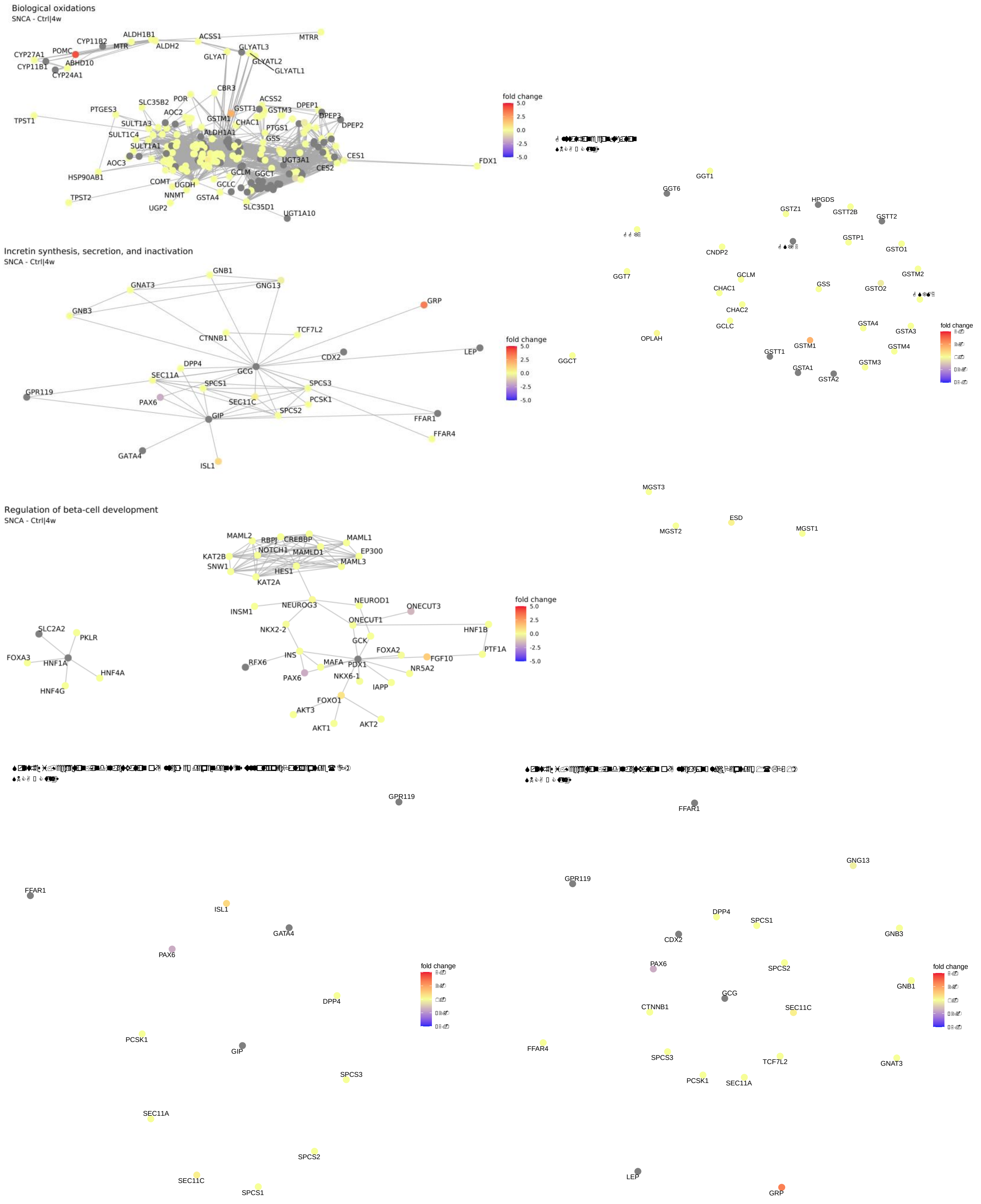

**Supplementary figure 2**

The six most differentially expressed pathways between SNCA and control mDA neurons are *i.* Biological oxidation, *ii.* Incretin synthesis, secretion and inactivation, *iii.* Glutathione conjugation, *iv.* Regulation of beta-cell development, *v.* Synthesis, secretion and inactivation of GIP and *vi.* Synthesis, secretion and inactivation of GLP-1

**Supplementary figure 3 – GLP-1R agonist reverses cellular pathology and prevents neurotoxicity in SNCA A53T and SNCAx3 midbrain dopaminergic neurons.**

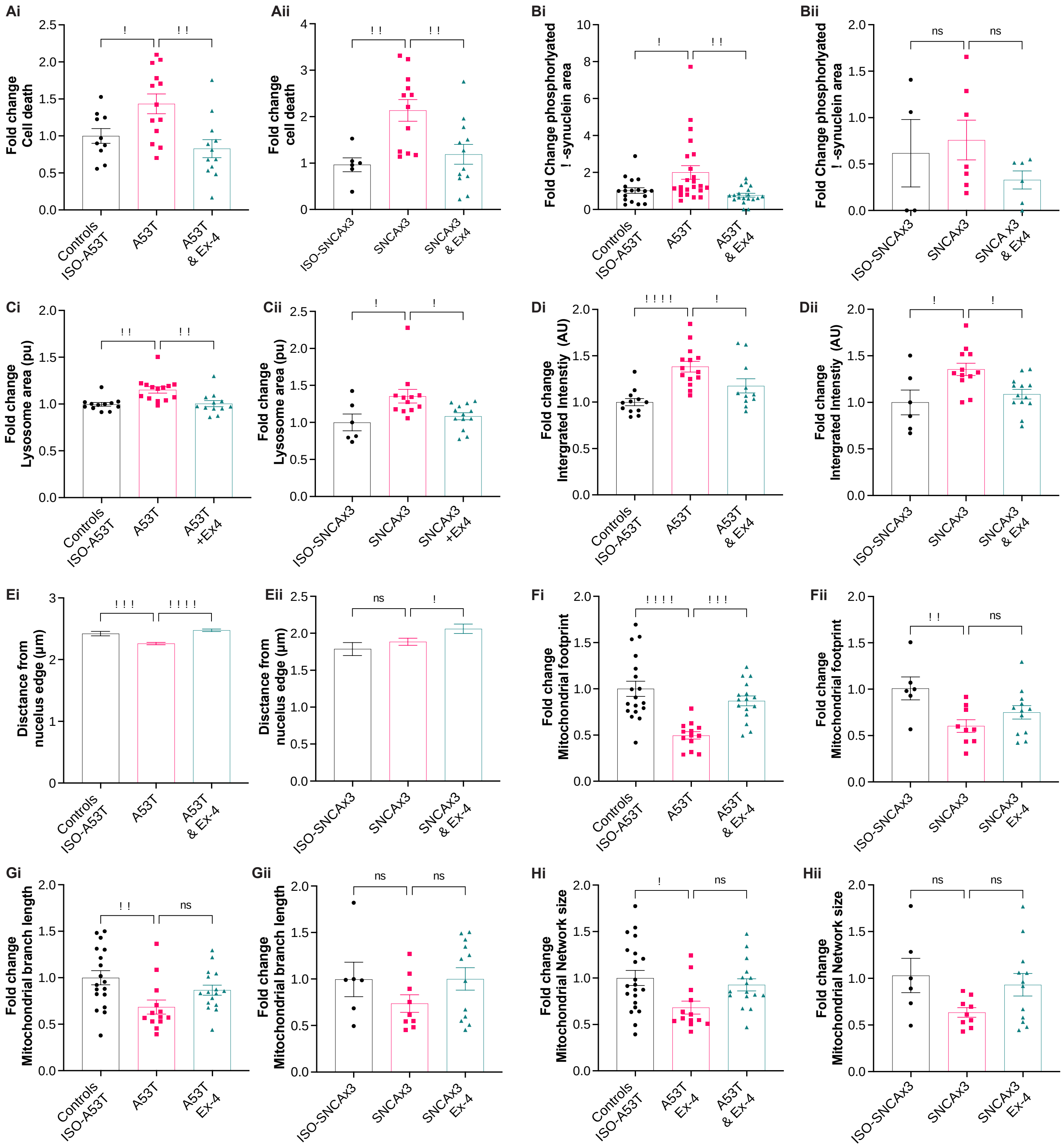

**Supplementary figure 3**

**A.** Fold change in cell death in A53T (**i**) and SNCAx3 (**ii**) mDA neurons ± Ex-4. **B.** Fold change in phosphorylated □-syn aggregates in A53T (**i**) and SNCAx3 (**ii**) mDA ± Ex-4. Fold change in lysosome area (**C**), integrated intensity (**D**) and distance from the nucleus edge (**E**) in A53T (**i**) and

SNCAx3 (**ii**) mDA neurons ± Ex-4. Fold change in mitochondria footprint (**F**), mitochondrial branch length (**G**) and network size (**H**) in A53T (**i**) and SNCAx3 (**ii**) mDA neurons ± Ex-4.

**Supplementary figure 4 - Proteomic expression alterations within MAPK signaling pathway and insulin receptor signaling pathway in post-mortem PD brain samples.**

A

| Protein ID | Substantia nigra | Parahippocampus | Frontal cortex |
|---|---|---|---|
| | Early vs Control | Early vs Control | Late vs Control |
| MAPK1 | 1.24 | 1.31 | -1.28 |
| MAP2K1 | 1.80 | 1.08 | 1.97 |

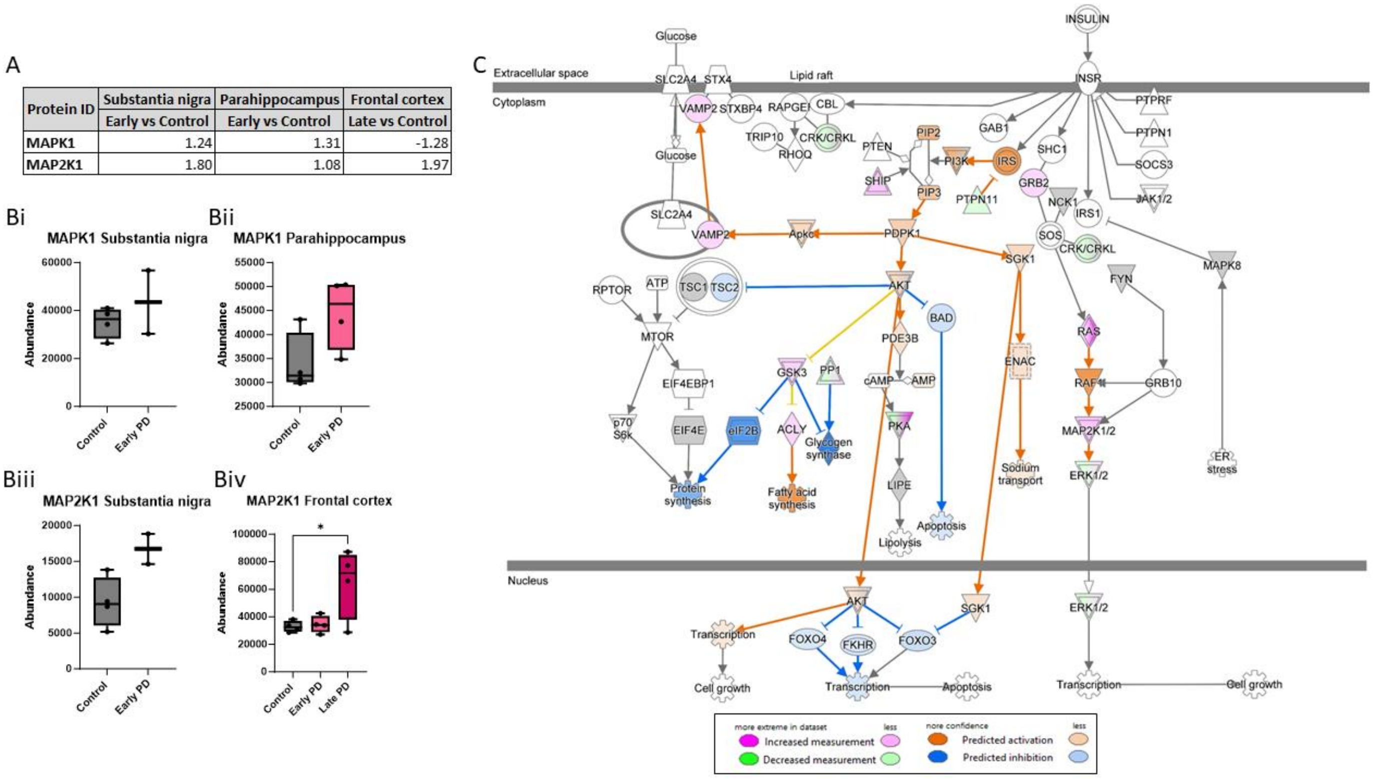


**Supplementary figure 4**

**A.** Table of fold change proteomic expression in substantia nigra, parahippocampus and frontal cortex brain regions in either early PD or late PD compared to controls for MAPK1 and MAP2K1. **B.** Abundance levels of MAPK1 (i and ii) and MAP2K1 (iii and iv) in each represented brain region: substantia nigra (i and iii), parahippocampus (ii) and frontal cortex (iv). Statistical significance measured by One-way ANOVA with Tukey multiple comparisons test *p < 0.05. **C.** Pathway diagram of insulin receptor signaling pathway from Qiagen IPA software with substantia nigra early PD vs control dataset overlaid. Pink represents any proteins upregulated in the dataset within the pathway and green represents any proteins downregulated. Predictions of activation (orange) or inhibition (blue) for the rest of the pathway based on these expression changes observed are then made by the IPA software

### Supplementary figure 5 - Clustering analysis for patients treated with exenatide (n=29)

A

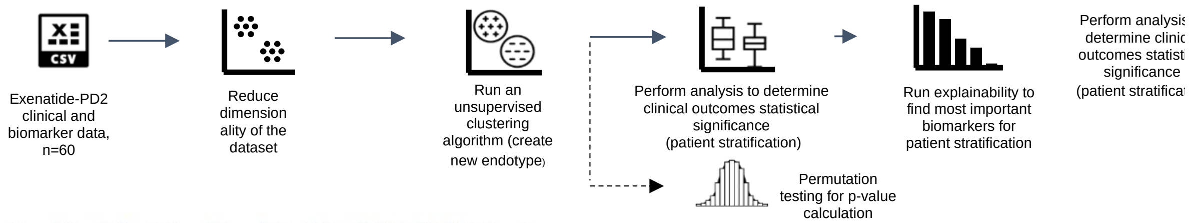


B

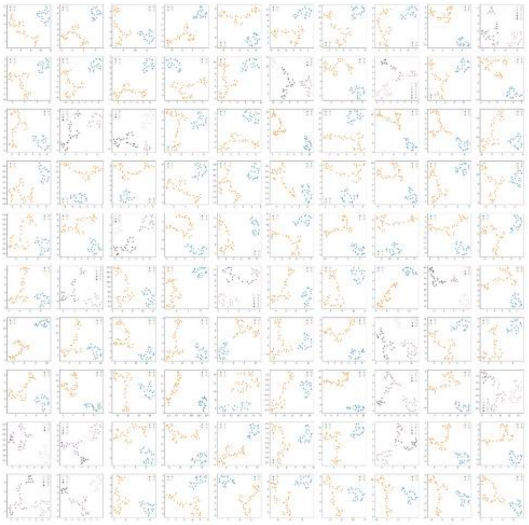

C

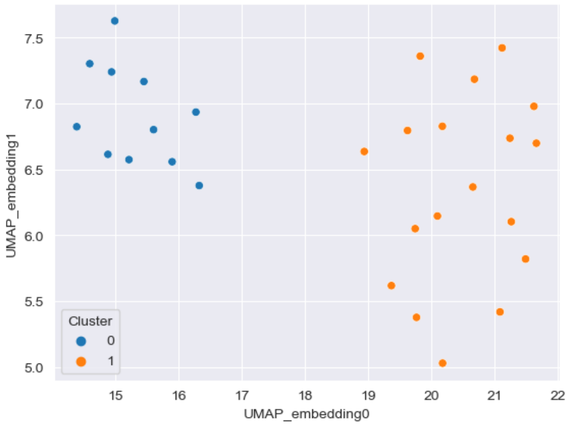


D

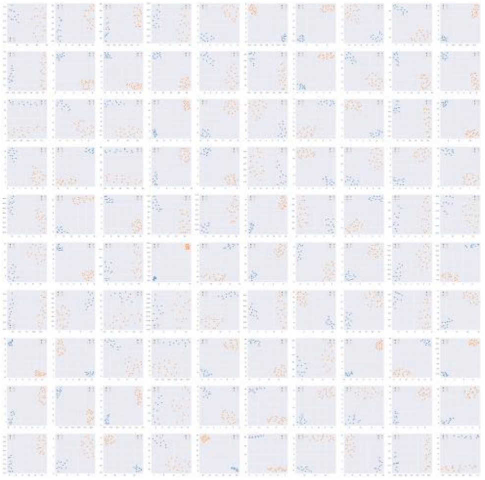

E

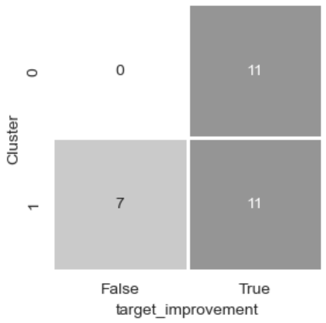


F

| Cluster | TS6K | TJNK | TP38 | PP38 | TERK | PJNK | PGSK-3B | TGSK-3B | PERK | Pan Tyr IRS | PMTOR | TMTOR | TIRS | IRS-312 | IRS-1616 | PAKT | TAKT |
|---|---|---|---|---|---|---|---|---|---|---|---|---|---|---|---|---|---|
| 0 | 6.55 | 6.15 | 6.73 | 6.01 | 6.71 | 6.39 | 5.16 | 5.3 | 5.66 | 4.8 | 5.92 | 5.78 | 8.02 | 9.64 | 8.23 | -0.42 | -0.02 |
| 0 | 6.47 | 5.91 | 6.22 | 5.79 | 6 | 5.83 | 4.79 | 5.09 | 5.66 | 4.83 | 6.73 | 6.69 | 7.05 | 9.53 | 7.04 | -1.08 | 0.34 |
| 0 | 6.22 | 5.54 | 5.9 | 5.99 | 6.06 | 6.06 | 4.55 | 4.75 | 5.65 | 5.24 | 7.01 | 6.91 | 7.28 | 9.33 | 7.2 | -0.71 | 0.71 |
| 0 | 6.46 | 5.72 | 6.2 | 6.05 | 6.38 | 6.17 | 4.92 | 5.02 | 5.76 | 4.74 | 6.39 | 5.92 | 7.23 | 6.61 | 7.93 | -1.61 | -0.12 |
| 0 | 6.73 | 6.43 | 6.45 | 6.12 | 6.3 | 6.19 | 5.09 | 5.33 | 5.86 | 5.96 | 6.42 | 6.2 | 7 | 8.87 | 7.7 | -1.02 | -0.22 |
| 0 | 6.84 | 6.31 | 6.51 | 6.35 | 6.45 | 6.58 | 5.51 | 5.24 | 6.28 | 7.84 | 6.8 | 6.81 | 8.31 | 9.96 | 8.79 | -0.71 | 1.38 |
| 0 | 5.91 | 5.61 | 6.1 | 5.58 | 6.23 | 5.65 | 4.58 | 4.91 | 5.61 | 5.48 | 6.3 | 6.26 | 6.86 | 8.57 |  | -0.78 | 0.96 |
| 0 | 6.36 | 5.88 | 6.53 | 6.95 | 6.5 | 6.11 | 4.67 | 4.96 | 5.83 | 5.09 | 6.69 | 6.56 | 7.02 | 8.89 | 7.57 | -1.24 | -0.22 |
| 0 | 6.27 | 5.63 | 6.26 | 5.67 | 6.35 | 5.68 | 4.71 | 4.93 | 5.52 | 4.89 | 6.44 | 6.3 | 7.3 | 9.7 | 8.11 | -0.39 | 1.44 |
| 0 | 6.19 | 5.85 | 6.51 | 6.73 | 6.59 | 5.87 | 4.68 | 5.02 | 5.6 | 5.09 | 5.63 | 5.4 | 6.83 | 8.81 | 7.25 | -0.43 | 3 |
| 0 | 6.26 | 5.99 | 6.2 | 5.89 | 6.45 | 5.89 | 4.84 | 5.04 | 6.17 | 4.88 | 6.26 | 6.09 | 8.27 | 9.62 | 8.16 | -1.14 | 0.6 |
| 1 | 5.74 | 5.24 | 5.86 | 5.06 | 6.11 | 5.21 | 4.38 | 5.04 | 4.92 | 5 | 4.95 | 4.64 | 6.66 | 9.03 | 7.66 | -0.82 | 0.12 |
| 1 | 5.98 | 5.29 | 5.95 | 5.11 | 5.96 | 5.21 | 4.37 | 4.63 | 4.91 | 4.93 | 5.48 | 5.64 | 6.7 | 8.3 | 7.11 | -1.24 | -0.2 |
| 1 | 5.49 | 5.44 | 5.96 | 5.45 | 5.89 | 5.28 | 4.16 | 4.54 | 5.01 | 4.69 | 5 | 5.06 | 5.67 | 7.34 | 6.81 | -1.08 | 0.42 |
| 1 | 5.47 | 5.18 | 5.96 | 4.75 | 5.92 | 4.91 | 4.32 | 4.65 | 4.87 | 4.73 | 5.67 | 5.67 | 6.7 | 9.13 | 7.57 | -0.89 | 0.26 |
| 1 | 5.9 | 5.63 | 6.05 | 5.42 | 6.01 | 5.53 | 4.01 | 4.33 | 5.23 | 5.02 | 6.16 | 6.19 | 7.27 | 9.08 | 7.58 | -0.65 | 0.85 |
| 1 | 5.29 | 5.09 | 6.04 | 5.15 | 6.08 | 5.4 | 3.97 | 4.91 | 5.21 | 4.61 | 5.78 | 5.51 | 6.39 | 8.35 | 7.7 | -0.76 | 1.59 |
| 1 | 5.86 | 5.28 | 5.74 | 6.06 | 5.86 | 6.1 | 4.74 | 4.93 | 5.89 | 4.74 | 5.47 | 5.37 | 5.89 | 7.55 | 6.64 | -0.97 | 0.74 |
| 1 | 5.82 | 5.37 | 5.78 | 5.34 | 6.11 | 5.42 | 4.84 | 5.02 | 5.28 | 4.34 | 5.79 | 5.8 | 5.23 | 7.95 | 6.81 | -1.56 | -0.82 |
| 1 | 5.62 | 4.93 | 5.38 | 4.71 | 5.67 | 5.02 | 4.01 | 4.34 | 4.93 | 4.57 | 6.24 | 6.11 | 9.27 | 8.52 | 7.82 | -1.35 | 0.03 |
| 1 | 5.86 | 5.42 | 5.88 | 5.26 | 6.11 | 5.32 | 4.43 | 4.77 | 5.71 | 4.57 | 5.44 | 5.19 | 6.02 | 8.42 | 8.2 | -0.94 | 1.37 |
| 1 | 5.9 | 5.05 | 5.39 | 5.58 | 5.65 | 5.84 | 4.38 | 4.55 | 5.78 | 4.84 | 6.3 | 5.95 | 6.29 | 8.91 | 7.32 | -0.89 | 0.34 |
| 1 | 5.36 | 5.42 | 5.89 | 4.79 | 5.89 | 4.93 | 4.06 | 4.55 | 4.8 | 4.62 | 5.09 | 5 | 5.5 | 6.92 |  | -1.97 | -0.89 |
| 1 | 5.93 | 5.52 | 6.1 | 5.46 | 6.25 | 5.61 | 4.32 | 4.65 | 5.4 | 4.91 | 6.45 | 6.31 | 5.89 | 8.19 | 7.49 | -1.31 | -0.63 |
| 1 | 5.89 | 5.42 | 6.03 | 5.4 | 5.94 | 5.48 | 4.19 | 4.67 | 5.38 | 4.72 | 6.2 | 5.95 | 6.99 | 8.87 | 7.27 | -0.62 | 1.24 |
| 1 | 5.86 | 5.32 | 5.98 | 5.33 | 5.95 | 5.46 | 4.2 | 4.75 | 5.18 | 4.43 | 5.9 | 5.15 | 5.91 | 7.27 | 7.93 | -0.94 | 0.96 |
| 1 | 5.73 | 5.25 | 5.94 | 5.01 | 6.07 | 5.1 | 4.29 | 4.66 | 5.06 | 4.74 | 6.32 | 6.14 | 6.98 | 9.38 | 6.94 | -0.99 | 0.42 |
| 1 | 5.72 | 5.05 | 5.42 | 4.97 | 5.67 | 5.09 | 4.41 | 4.6 | 4.84 | 4.54 | 5.97 | 5.81 | 6.5 | 8.46 | 7.09 | -1.66 | -0.4 |
| 1 | 5.66 | 5.48 | 6.03 | 5.04 | 5.92 | 4.91 | 4.79 | 4.62 | 4.91 | 4.68 | 5.27 | 5.34 | 6.34 | 8.86 |  | -1.27 | -0.25 |

**Supplementary figure 6**

**A**. Outline of methods used to generate predictive biomarkers. **B.** Clustering 100 times with different random initiations to ensure robustness of clusters – derived same 2 clusters 80% of the time. **C.** Clustering analysis using only the exenatide treated individuals. **D.** Clustering 100 times with different random initiations to ensure robustness of clusters – derived same 2 clusters 83% of the time. **E.** 100% of individuals in Cluster 0 improved with exenatide treatment, whereas only 60% of participants improved with exenatide treatment if in Cluster 1. **F.** Permutation testing to see which variables are significantly associated with the difference between cluster 0 and cluster 1 highlighted elevated levels of MAPK pathway kinases as predictors of response to exenatide.

**Supplementary Table 1 – hiPSC lines used in this study**

| hiPSC line | Mutation | Age of donor | Sex of donor | Source | RRID |
|---|---|---|---|---|---|
| No disease Control 1 | None | 78 | Male | Reprogrammed in house, Kunath lab EDi046-A | |
| No disease Control 2 | None | 51 | Male | Cedars Sinai iPSC Core Repository CS0002iCTR-nxx | |
| No disease Control 3 | None | Unknown | Female | Thermo Fisher Scientific A18945 | CVCL_RM92 |
| No disease Control 4 | None | 64 | Male | Coriell ND41866 | CVCL_Y838 |

| | | | | | |
|---|---|---|---|---|---|
| No disease Control 5 | None | 60-64 | Female | EBiSC<br>WTSli017-B<br>HPSI0114i-lexy_1 | CVCL_AE31 |
| No disease Control 6 | None | 55-59 | Male | EBiSC<br>WTSli018-A<br>HPSI0114i-kolf_3 | CVCL_AE30 |
| No disease Control 7 | None | 55-59 | Male | EBiSC<br>WTSli019-B<br>HPSI0114i-iisa_1 | CVCL_AE22 |
| A53T 1 & isogenic control | *SNCA* A53T | 54 | Female | StemBANCC<br>SFC828 (STBCi019-A) | CVCL_RB71 |
| A53T 2 | *SNCA* A53T | 57 | Male | StemBANCC<br>SFC829 (STBCi023-C) | CVCL_RB78 |
| *SNCA* x3 | *SNCA* locus x3 | Unknown | Female | StemBANCC<br>SFC831 (STBCi024-C) | CVCL_RB81 |
| *SNCA* x3 & isogenic control | *SNCA* locus x3 | Unknown | Female | As reported [54]<br>EDi001-B | CVCL_ZA47 |

**Supplementary Table 2 – List of antibodies used in this study**

| Protein | Company | Catalogue | RRID | Species | Dilution |
|---|---|---|---|---|---|
| TH | Abcam | ab137869 | AB_2801410 | Rabbit | 1:500 (ICC) |

| | | | | | |
|---|---|---|---|---|---|
| | | | | | 1:200 (Flow) |
| TH | Abcam | ab76442 | AB_1524535 | Chicken | 1:500 |
| MAP2 | Abcam | ab183830 | AB_2895301 | Rabbit | 1:500 |
| GFAP | Abcam | ab4674 | AB_304558 | Chicken | 1:250 |
| Total alpha-synuclein | Abcam | ab138501 | AB_2537217 | Rabbit | 1:200 |
| Filament alpha-synuclein | Abcam | ab209538 | AB_2714215 | Rabbit | 1:50-1:100 |
| Phosphorylated alpha-synuclein | Abcam | ab51253 | AB_869973 | Rabbit | 1:1000 |
| Goat pAb anti-Mouse IgG Alexa Fluor 555 | Abcam | ab150114 | AB_2687594 | Goat | 1:500 |
| Goat pAb anti-Rabbit IgG Alexa Fluor 647 | Abcam | ab150079 | AB_2722623 | Goat | 1:500 |
| Goat pAb anti-Chicken IgG Alexa Fluor 488 | Abcam | ab150169 | AB_2636803 | Goat | 1:500 |

**Supplementary Table 3 – List of protocols used in this study.**

| Protocol | DOI |
|---|---|
| Human iPSC cell culture | dx.doi.org/10.17504/protocols.io.81wgby7dnvpk/v1 |
| Generation of midbrain dopaminergic neurons | dx.doi.org/10.17504/protocols.io.x54v9j7ezg3e/v1 |
| Generation of cortical neurons | 10.1038/nprot.2012.116 |
| Generation of astrocytes | dx.doi.org/10.17504/protocols.io.8epv5xmmng1b/v1 |
| Single-cell RNA-seq | dx.doi.org/10.17504/protocols.io.6qpvr4dxpgmk/v1 |
| Immunocystochemistry | dx.doi.org/10.17504/protocols.io.q26g74w79gwz/v1 |
| Sample preparation and ELISA | dx.doi.org/10.17504/protocols.io.dm6gp35jdvzp/v1 |
| SAVE imaging | dx.doi.org/10.17504/protocols.io.n92ldmqkol5b/v1 |
| SiMPull | dx.doi.org/10.17504/protocols.io.36wgq3bz3lk5/v1 |
| Quantitative polymerase chain reaction (qPCR) | dx.doi.org/10.17504/protocols.io.36wgqj5rkvk5/v1 |
| Multiplex live imaging | dx.doi.org/10.17504/protocols.io.8epv5xdq5g1b/v1 |
